# TIPRL1 and its ATM-dependent phosphorylation promote radiotherapy resistance in head and neck cancer

**DOI:** 10.1101/2023.08.30.555468

**Authors:** Célie Cokelaere, Rüveyda Dok, Emanuela E. Cortesi, Peihua Zhao, Anna Sablina, Sandra Nuyts, Rita Derua, Veerle Janssens

## Abstract

TIPRL1 (target of rapamycin signaling pathway regulator-like 1) is a known interactor and inhibitor of protein phosphatases PP2A, PP4 and PP6 – all pleiotropic modulators of the DNA Damage Response (DDR). Here, we describe a new role for TIPRL1 in the radiotherapy (RT) response of Head and Neck Squamous Cell Carcinoma (HNSCC). TIPRL1 expression was found increased in tumor versus non- tumor tissue, with high tumoral TIPRL1 expression associating with lower locoregional control and decreased survival of RT-treated patients. TIPRL1 deletion in HNSCC cells resulted in increased RT sensitivity, a faster but prolonged cell cycle arrest, increased micronuclei formation and an altered proteome-wide DDR. Upon irradiation, ATM phosphorylates TIPRL1 at Ser265, contributing to TIPRL1-mediated RT resistance. Mass spectrometry analysis identified DNA-PKcs, RAD51 and nucleosomal histones as novel TIPRL1 interactors. Histone binding, although stimulated by RT, was adversely affected by TIPRL1 Ser265 phosphorylation. Our findings underscore a clinically relevant role for TIPRL1 and its ATM-dependent phosphorylation in RT resistance through modulation of DNA damage checkpoint activation and repair.

## Introduction

Therapy resistance remains one of the major hurdles in cancer treatment. Head and neck squamous cell carcinoma (HNSCC) is the sixth most frequently occurring cancer worldwide (Bray et al, 2018), and is predominantly treated with radiotherapy (RT). HNSCC tumors arise in the oral cavity, larynx and pharynx, and are subdivided into two etiology groups. Human papillomavirus-positive (HPV+) HNSCC show better RT response rates, and HPV positivity is considered as an independent prognostic marker. The etiology of HPV-negative (HPV-) HNSCC is linked to tobacco and/or alcohol use, and these tumors are genetically more heterogeneous and show significantly higher RT resistance (Ang et al, 2010; Lassen, 2010; Leemans et al, 2011; Suh et al, 2014; Vigneswaran & Williams, 2014). Despite technical advances in surgery and radiotherapy, up to 50% of treated HNSCC patients diagnosed with a locally advanced tumor develop locoregional recurrences, second primary tumors or distant metastases due to therapy resistance, with minimal residual options for re-irradiation or salvage surgery (Leemans et al, 2011; Suh et al, 2014). These dismal numbers emphasize the need for new therapeutic strategies sensitizing HNSCC for RT, and for better patient stratification to improve overall survival (Buglione et al, 2017; Magrini et al, 2016).

Protein Phosphatase 2A (PP2A) represents a large family of Serine (Ser)/Threonine (Thr) phosphatases mainly possessing tumor suppressive functions (Eichhorn et al, 2009; Janssens & Goris, 2001; Meeusen & Janssens, 2018). The majority of PP2A complexes are trimers consisting of a catalytic C, a scaffolding A and a regulatory B subunit, of which the latter determines substrate specificity (Lambrecht et al, 2013). PP2A complexes play an important role in the DNA damage response (DDR), largely acting to prevent DDR activation in cells devoid of DNA damage, or to cease the DDR when damage has been repaired (Freeman & Monteiro, 2010; Lee & Chowdhury, 2011; Peng & Maller, 2010; Sule et al, 2022; Zheng et al, 2015). As PP2A’s functions are mainly tumor suppressive, PP2A is frequently inactivated in many cancer types (Haanen et al, 2022; Sangodkar et al, 2016; Velmurugan et al, 2018; Wang et al, 2019). In HNSCC, PP2A inactivation largely occurs in a non-genomic way, by overexpression of specific cellular PP2A inhibitors (Gouttia et al, 2022; Junttila et al, 2007; Katz et al, 2010; Leopoldino et al, 2012; Patel et al, 2008; Ventelä et al, 2014). Like that, overexpression of CIP2A (Cancerous inhibitor of PP2A) or SET (Suvar/Enhancer of zeste/Trithorax) correlates with poor patient survival, enhanced HNSCC cell survival and decreased cisplatin response (Alzahrani et al, 2020; Böckelman et al, 2011; Leopoldino et al, 2012; Liu et al, 2014; Ouchida et al, 2019; Routila et al, 2016; Sobral et al, 2014). However, whether other PP2A inactivating mechanisms could be of clinical relevance in HNSCC etiology or therapy response, remains poorly understood.

Here, we focused on TIPRL1 (target of rapamycin (TOR) signaling pathway regulator-like 1) as another presumed cellular PP2A inhibitor (Feng et al, 2016; Gingras et al, 2005; McConnell et al, 2007; Rosales et al, 2015). Originally discovered in yeast as a regulator of TOR signaling (Jacinto et al, 2001), TIPRL1 was subsequently identified as a binding partner of PP2A, PP4 and PP6 phosphatases (Gingras et al, 2005). *In vitro*, TIPRL1 inhibits the activity of the PP2A AC dimer, but not of trimeric PP2A B56 complexes (Haesen et al, 2016; McConnell et al, 2007; Smetana & Zanchin, 2007). Structural data revealed TIPRL1 binding to the C-terminal region of PP2A C (Scorsato et al, 2016; Smetana & Zanchin, 2007; Wu et al, 2017) requiring integrity and demethylation of its carboxyterminal tail (Scorsato et al, 2016; Wu et al, 2017), suggesting that TIPRL1-mediated PP2A inhibition is limited to unmethylated PP2A complexes (Wu et al, 2017). Whether similar rules apply to TIPRL1 inhibition of PP4 and PP6 complexes is unclear, although for PP4, TIPRL1 interaction with a specific trimer (PP4C-R2-R3) has been demonstrated (Gingras et al, 2005). Besides its phosphatase inhibitory role, TIPRL1 may also play a role in PP2A C recycling and reassembly of PP2A holoenzymes (Sents et al, 2013; Wu et al, 2017), occurring during PP2A holoenzyme biogenesis or following holoenzyme disassembly in stressed cells (Kong et al, 2009). For this function, TIPRL1 may collaborate with alpha4 (*IGBP1*), another common binding partner and regulator of PP2A/PP4/PP6 (Smetana & Zanchin, 2007; Wu et al, 2017). This dual role in phosphatase regulation may also explain several seemingly contradictory reports on the poorly understood cellular functions of TIPRL1 – with some papers consistent with its phosphatase inhibitory role (McConnell et al, 2007; Nakashima et al, 2013; Rosales et al, 2015), and others rather underscoring the opposite (Feng et al, 2016; Gingras et al, 2005; Jacinto et al, 2001; Song et al, 2012).

In this study, we identified TIPRL1 as a novel substrate of Ataxia Telangiectasia Mutated (ATM) kinase in irradiated HPV- HNSCC cells, and found that TIPRL1 downregulation or expression of a non-phospho TIPRL1 mutant sensitized the cells for RT. RAD51, DNA-PKcs and histones were identified as novel TIPRL1 interactors. TIPRL1 deletion dysregulated the DDR at multiple levels, eventually resulting in a faster cell cycle arrest but decreased DNA repair. Underscoring the potential clinical relevance of these findings, TIPRL1 expression was found increased in a significant fraction of HNSCC tumors, which correlated with decreased locoregional control and decreased patient survival. Thus, TIPRL1 could be useful as a new predictive marker and target for HPV- HNSCC RT response.

## Results

### High TIPRL1 expression in HNSCC tumors correlates with decreased locoregional control and disease-free survival in RT-treated patients

Currently, the status of TIPRL1 expression in HNSCC is unknown. Upon examination of the HNSCC TCGA (Firehose Legacy) study (n=563) through the cBioportal database (Cerami et al, 2012; Gao et al, 2013; The Cancer Genome Atlas Network, 2015), we found significantly increased TIPRL1 expression in HNSCC tumor tissues (n=519) compared to normal epithelial tissues (n=44) (t-test: p<0.0001) (**Fig 1A**). Moreover, HPV- HNSCC patients with high TIPRL1 expression who received RT showed a trend for lower disease-free survival than those with low TIPRL1 expression (**Fig 1B, left panel**). Notably, this trend was not observed for the patients who received treatments other than RT (**Fig 1B, right panel**).

**Figure 1.**
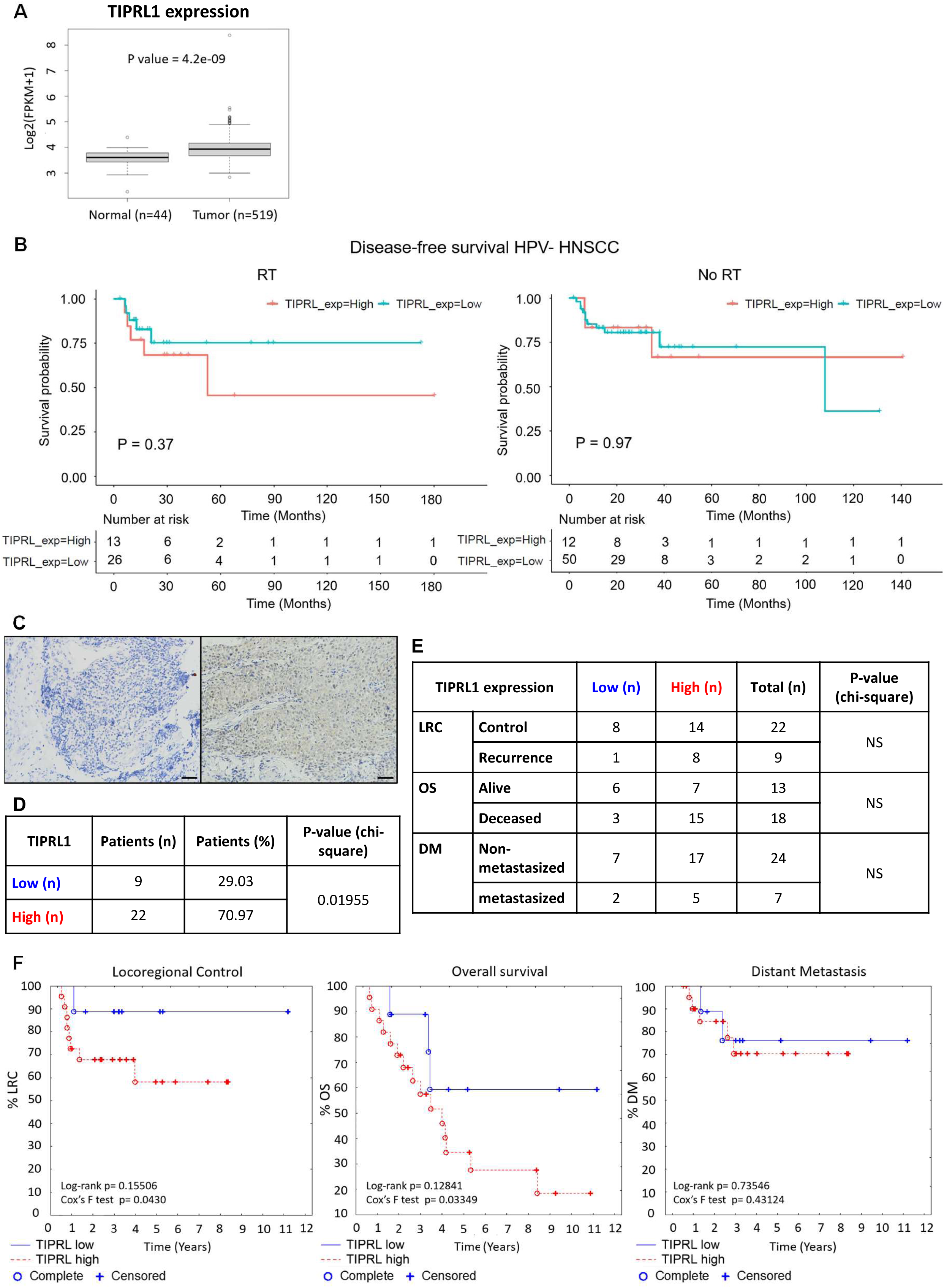
TIPRL1 expression in HNSCC patient samples and its predictive role in HPV- HNSCC patients treated with RT. **A** Examination of TIPRL1 expression in 563 samples (519 tumor, 44 non-tumor tissues) of the TCGA HNSCC Firehose Legacy study (The Cancer Genome Atlas Network, 2015), indicating increased expression in tumor versus non-tumor tissue (t-test: p<0.0001). **B** Using data from the same study, patients with HPV- HNSCC were divided in a group who received RT (left panel) and a group who did not (right panel), and further stratified into TIPRL1-high (above mean) and TIPRL1-low (below mean) tumors based on the average TIPRL1 expression level in all tumors. Patients with tumors expressing high levels of TIPRL1 and treated with RT showed a lower disease-free survival (DFS) than those with low TIPRL1 expression (left panel). This trend cannot be seen for patients that received other treatment than irradiation (right panel) (log-rank tests). **C** Representative image of low TIPRL1 (left panel) or high TIPRL1 (right panel) expression in HNSCC samples from an independent UZ Leuven patient cohort, as visualized by TIPRL1 immunohistochemical staining. Bar: 50 µm. **D** Chi-square test was used to determine difference in number of tumors with low or high TIPRL1 expression (31 HNSCC patients). **E** Chi-square test was performed to evaluate whether patients with low or high TIPRL1 expression show differences for Locoregional Control (LRC), Overall Survival (OS) or Distant Metastasis (DM). NS: not significant (p>0.05). **F** Kaplan-Meier curves (n=31) show a trend (log-rank test) or significance (Cox’s F test) for lower LRC and OS for patients with high TIPRL1 expression. No significant differences for DM (log-rank and Cox’s F tests: p>0.05).

To independently confirm this observation, we performed immunohistochemical analysis for TIPRL1 expression on 31 pretreatment HNSCC tumor biopsies collected at the University Hospitals Leuven (**Fig 1C).** TIPRL1 expression, seen both in the nucleus and the cytoplasm of cancer and non-cancer cells, was significantly increased in 71% of samples (22/31, χ^2^ test: p<0.02) (**Fig 1D**). Although the observed frequencies of locoregional control (LRC), overall survival (OS) or distant metastasis (DM) did not differ between patients with high or low TIPRL1 expression (χ^2^ test: non-significant) (**Fig 1E**), the Kaplan-Meier curves showed a borderline significant decrease in LRC (log-rank test: p=0.16; Cox’s F test: p<0.05) and OS (log-rank test: p=0.13; Cox’s F test: p<0.05) for RT-treated patients with high TIPRL1 expression, while no differences were found for occurrence of DM (log-rank test: p=0.74; Cox’s F test: p=0.43) (**Fig 1F**). These findings align with TCGA results (**Fig 1B**) and indicate that, as reported for CIP2A (Junttila et al, 2007; Katz et al, 2010) and SET (Leopoldino et al, 2012) PP2A inhibitors, TIPRL1 is overexpressed in a high percentage of HNSCC tumors, correlating with lower OS and decreased LRC. These data thus imply a predictive role for TIPRL1 expression in HPV- HNSCC patients treated with radiotherapy.

### *TIPRL* deletion via CRISPR/Cas9 editing increases RT sensitivity of HPV- SQD9 cells

To identify a suitable cell model for subsequent mechanistic studies, we determined TIPRL1 expression in four HPV- HNSCC cell lines (Cal27, SQD9, SC263 and SCC61) and the primary human tonsil epithelial cell line (HTEpiC) by immunoblotting (**Fig 2A**). An overexpression of TIPRL1 was seen in all four cancer cell lines, although to different extents (**Fig 2A**). As TIPRL1 expression was highest in the RT-resistant SQD9 cell line (Smeets et al, 1994), SQD9 cells were selected as our subsequent cell model.

**Figure 2.**
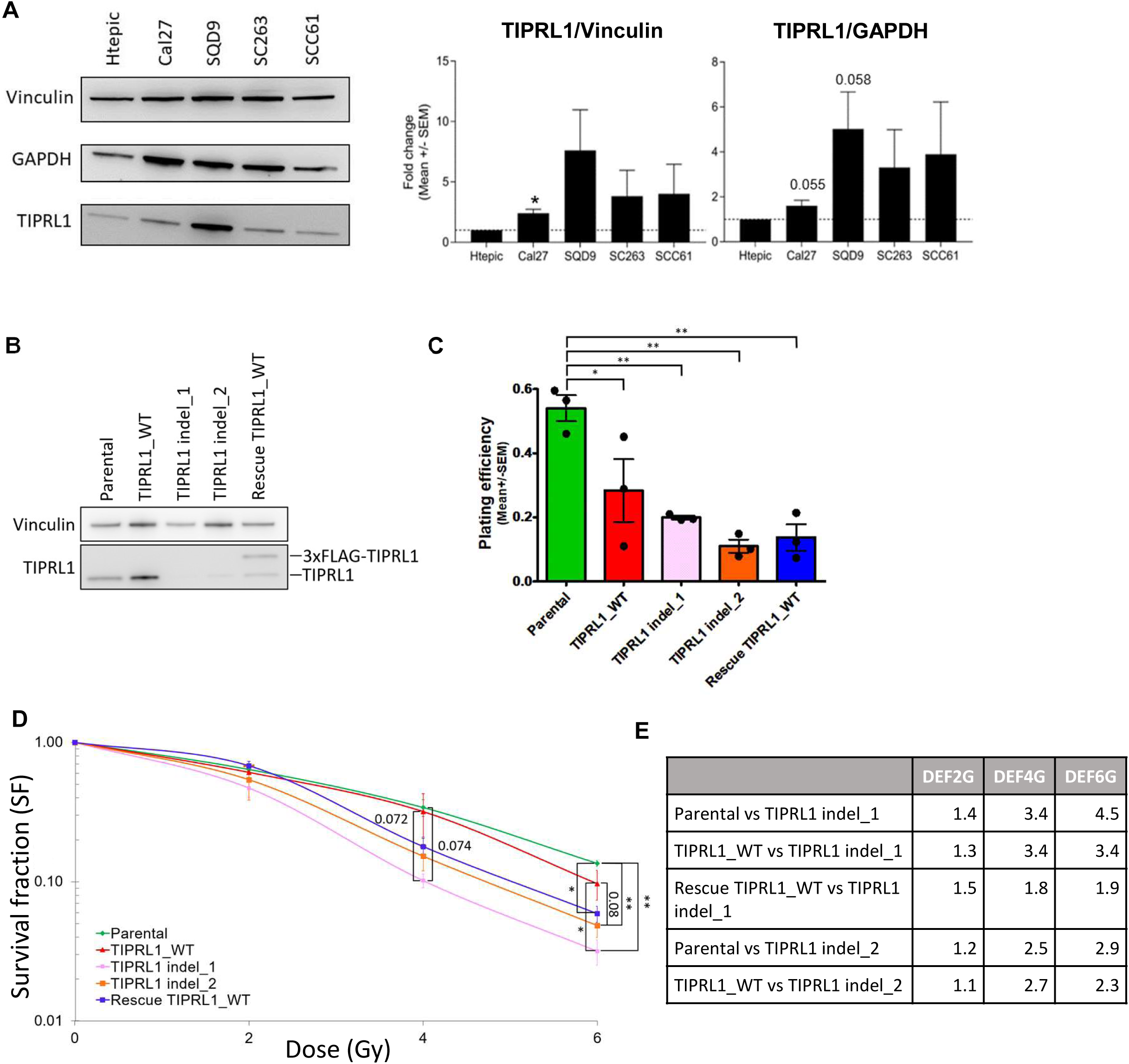
TIPRL1 expression in HPV- HNSCC cell lines and the radiosensitizing effect of *TIPRL* indel mutations in SQD9 cells. **A** Western blot analysis of total protein lysates of Htepic (non-cancerous), Cal27, SQD9, SC263 and SCC61 HNSCC cells to monitor relative TIPRL1 expression (representative blot). Two housekeeping genes (vinculin and GAPDH) were used as loading controls for quantification (n=6, one sample t-test, *: p<0.05). **B** Western blot analysis of TIPRL1 expression in the selected CRISPR clones and rescue TIPRL1 cell line. As the start codon of the TIPRL1 coding sequence was not removed within the 3xFLAG-TIPRL1 rescue construct, some untagged TIPRL1 is also re-expressed in the rescue TIPRL1 cell line. Vinculin was used as loading control (n=1). **C** Plating efficiency was expressed as the ratio between the number of colonies and the number of cells seeded (n=3, mean ± SEM, one-way ANOVA followed by Tukey’s multiple comparison test, *: p<0.05, **: p<0.01). **D** Survival fractions (SF, logarithmic scale) after different doses of RT (0, 2, 4 and 6 Gy) of the different cell lines are shown as the mean clonogenic survival (n=3, mean ± SEM, one-way ANOVA followed by Tukey’s multiple comparison test, *: p<0.05, **: p<0.01). **E** Dose Enhancement factors (DEF) were calculated for different doses of RT (2, 4 and 6 Gy) by making the ratios of the survival fractions of TIPRL1 expressing cells vs *TIPRL* indel cells (n=3).

To assess the potential role of TIPRL1 in the RT response of SQD9 cells, we introduced indel mutations in *TIPRL* using the CRISPR/Cas9 technique. With the PAM sequence of our guide RNA located in intron 4 of *TIPRL* (**gRNA1, Appendix Table S1**), we were able to induce an early stop codon in exon 5, giving rise to *TIPRL* indel SQD9 cell clones specifically targeting TIPRL1, but not the shorter non-PP2A binding TIPRL2 isoform (McConnell et al, 2007). As our negative control cell line, we selected a clone that underwent the CRISPR technique but maintained comparable TIPRL1 protein expression as the parental SQD9 cells (further referred to as TIPRL1_WT). Two *TIPRL* indel clones (TIPRL1 indel_1 and TIPRL1 indel_2), in which the indel mutation was verified by genomic sequencing (**Appendix Fig S1A**), were used for further experiments. While in silico analysis predicted the *ZNF317* gene as a potential off-target of gRNA1, Sanger sequencing did not detect any *ZNF317* mutations in either indel clone (**Appendix Fig S1B**). Immunoblotting analysis further confirmed absence of TIPRL1 protein expression in both *TIPRL* indel clones **(Fig 2B)**. We also stably reintroduced 3xFLAG-tagged TIPRL1 WT in TIPRL1 indel_1, to generate a rescued cell line (Rescue TIPRL1_WT). SQD9 parental, TIPRL1_WT and Rescue TIPRL1_WT all showed comparable TIPRL1 expression levels (**Fig 2B**).

To examine the effect of TIPRL1 expression on the RT response of SQD9 cells, 2D colony growth assays were executed following irradiation of the cells with increasing doses of RT up to 6 Gy. As CRISPR-based gene editing and lentiviral transduction are stressful events for the cells, we noticed that in the absence of RT the clonogenicity in TIPRL1_WT had diminished compared to the parental cell line and this was even further decreased in the rescued cell line (**Fig 2C**). Upon irradiation, lower survival fractions for both TIPRL1 indel_1 and TIPRL1 indel_2 were found compared to TIPRL1_WT and Parental cells (**Fig 2D**), suggesting that TIPRL1 expression confers increased resistance to RT. Moreover, re-expression of TIPRL1 (Rescue TIPRL1_WT) partially rescued the radiosensitive phenotype of the TIPRL1 indel_1 cell line (**Fig 2D**). Comparisons of the average dose enhancement factors (DEFs) further confirmed the radiosensitizing effects of *TIPRL* indel mutations (**Fig 2E**).

### SQD9 *TIPRL* indel cells show a faster and more prolonged checkpoint activation, cell cycle arrest, and increased micronuclei formation upon DNA damage

To determine whether decreased colony formation in irradiated *TIPRL* indel cells correlated with lower survival, we used a flow cytometry-based approach to compare the level of cell death in TIPRL1 indel_2 versus TIPRL1_WT cells upon irradiation. We identified minimal presence of Annexin V or propidium iodide-positive cells in both TIPRL1_WT and TIPRL1 indel_2 cells at 72h or 96h after 6 Gy irradiation (**Fig 3A, 3B**), suggestive for a cytostatic rather than a cytotoxic effect of RT in these cells. This hypothesis was further confirmed by flow cytometry following PI and BrdU staining (**Fig 3C, 3D; Fig EV1A, EV1B**). In non-irradiated cells, TIPRL1 deletion did not show any major effects on cell cycle distribution, except at the first time point (2h), when fewer cells were found in the G1 phase and more in S- and G2/M phase compared to TIPRL1_WT cells (**Fig 3C**), suggestive for some replicative difficulties early after plating. However, at later time points (4h up tot 24h), this problem was completely resolved (**Fig 3C**). In contrast, upon irradiation (6 Gy), this difference was maintained and became further pronounced, as evidenced by an increased number of TIPRL1 indel_2 cells in the S- and/or G2/M phases at 4h-24h post-RT (concomitantly with a decreased number of cells in the G1 phase at 2h-6h post-RT) as compared to TIPRL1_WT cells (**Fig 3D, Fig EV1A**). In accordance, there is a trend for a larger fraction of BrdU-positive TIPRL1_indel_2 cells at 6h and 24h post-RT than for the BrdU-positive TIPRL1_WT fraction at those time points (**Fig EV1B**). Hence, the loss of TIPRL1 provoked an earlier cell cycle arrest and an accumulation in S and G2/M phase upon RT. Consistently, we found increased RT-induced Ser345 phosphorylation of Chk1 in *TIPRL* indel cells at 1h and 24h post-RT when compared with Parental SQD9 cells and WT TIPRL1 rescued cells (**Fig 3E, 3F**) – indicative for a more pronounced (1h) and prolonged (24h) Chk1-induced checkpoint activation in irradiated TIPRL1-depleted cells.

**Figure 3.**
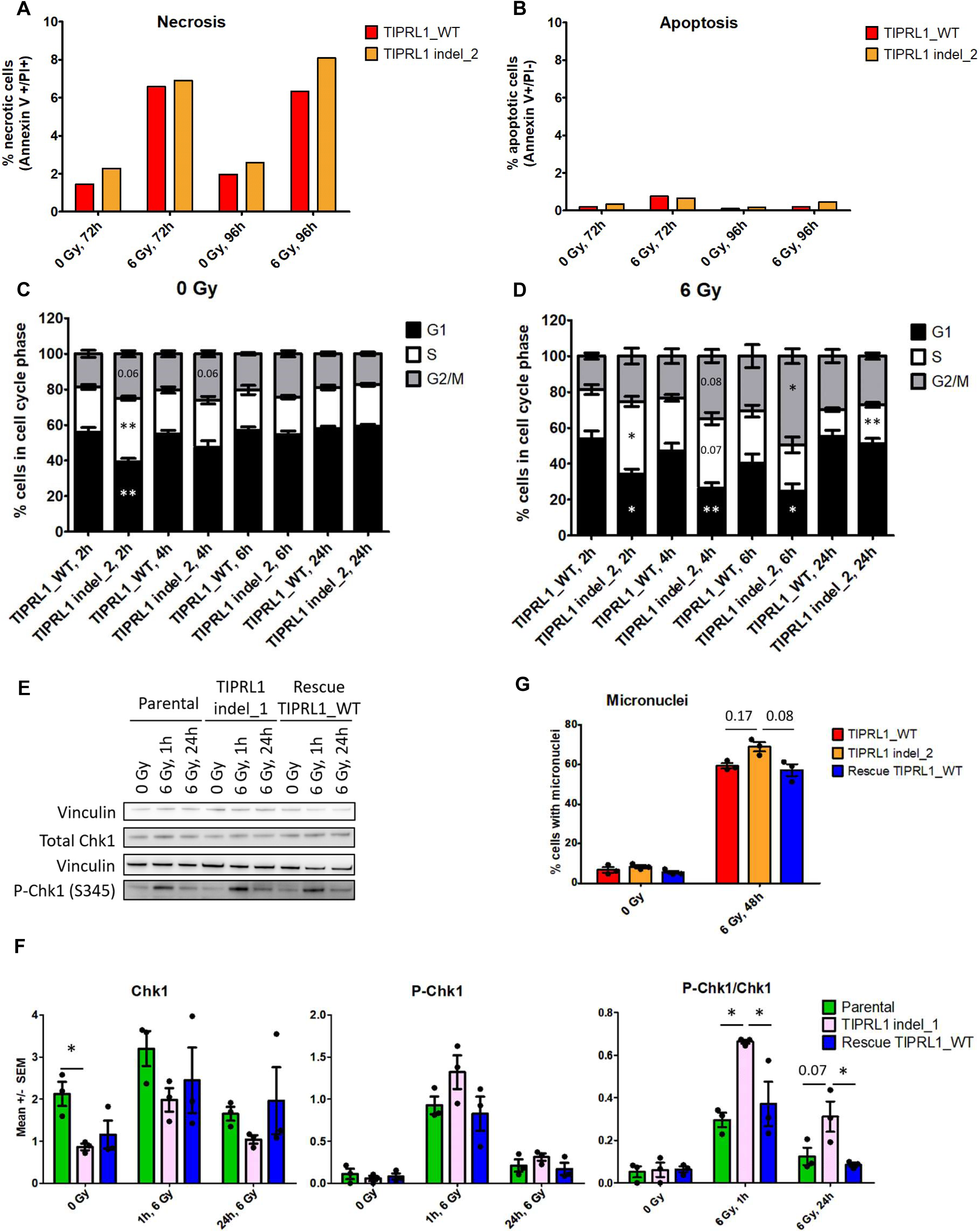
TIP*RL* indel mutations result in more micronuclei formation, a faster and prolonged S phase checkpoint arrest after DNA damage. **A, B** Flow cytometry using PI and Annexin V staining was executed 72h and 96h after treatment with 6 Gy of SQD9 cells with or without a *TIPRL* indel mutation (n=1). Percentage Annexin V and PI positive, or necrotic cells (**A**); Percentage Annexin V positive and PI negative, or apoptotic cells (**B**). **C, D** Cell cycle distribution using PI and BrdU staining for indicated cell lines without (**C**) or with 6 Gy irradiation (**D**). T-test was used to determine differences in percentage of cells in different cell cycle phases between TIPRL1_WT or TIPRL1 indel_2 cells at a certain time point with or without irradiation (n=3, *: p<0.05, **: p<0.01). **E, F** Western blot analysis of total protein lysates of (irradiated) SQD9 Parental, TIPRL1 indel_1 and rescue TIPRL1_WT cells (representative blot) (**E**), and corresponding quantification (**F**). Western blots were developed with primary Chk1 and p-Chk1 (Ser345) antibodies. Vinculin was used as loading control (n=3, Mean ± SEM, one-way ANOVA followed by Tukey’s multiple comparison test, *: p<0.05). **G** SQD9 cells expressing TIPRL1 WT or harboring *TIPRL* indel mutations were treated with 6 Gy irradiation. DAPI staining was performed 48h after irradiation to assess the percentage of micronucleated cells (n=3, mean ± SEM, one-way ANOVA followed by Tukey’s multiple comparison test).

The slower cell cycle progression of irradiated TIPRL1-depleted cells could be suggestive for impaired DNA-damage repair. Hence, we first quantified γ-H2AX foci in the irradiated cells at different times post-RT. While no differences were observed in γ-H2AX foci number between TIPRL1 WT or indel cells 1h-3h post-RT (**Fig EV1C**), the TIPRL1 indel_1 cells showed an increased number of foci compared to TIPRL1_WT cells 24h post-RT (**Fig EV1C**). However, because this phenotype could not be clearly rescued by TIPRL1 re-expression and was not observed in the other TIPRL1 indel clone (TIPRL1 indel_2), we cannot make a strong conclusion about a potential delay in repair of γ-H2AX-associated DNA lesions. Therefore, we also assessed micronuclei formation 48h after irradiation (6 Gy). An increase in micronuclei number was observed in *TIPRL* indel cells compared to TIPRL1_WT control cells, which could be reverted by TIPRL1 re-expression in the Rescue TIPRL1_WT cells (**Fig 3G**). This suggests that *TIPRL* indel cells indeed maintain more DNA damage after irradiation.

### Deletion of TIPRL1 affects the global HNSCC cell proteome after radiotherapy

We next used unbiased LC-MS/MS to assess potential changes in the total proteome of TIPRL1_WT versus TIPRL1 indel_2 cells, under basal conditions and at 4h, 24h and 48h post-irradiation. Given the known role of TIPRL1 as a regulator of PP2A-like phosphatases, we first assessed potential expression differences in PP2A, PP4 and PP6 subunits and regulators between TIPRL1_WT and TIPRL1 indel_2 cells under basal and irradiated conditions (**Fig 4A**). When we compared normalized protein abundances of all identified PP2A-like subunits/regulators (13 proteins) as a function of time after RT, it became apparent that higher expression levels were consistently present in the TIPRL1 indel_2 cells in any given condition relative to TIPRL1_WT control (**Fig 4A**). This trend was independently validated by immunoblotting for PP2A C, PP2A B55α and PP6C (**Fig 4B-4D**). Thus, these data seem to confirm a role for TIPRL1 in overall expression/stability regulation of PP2A-like phosphatases.

**Figure 4.**
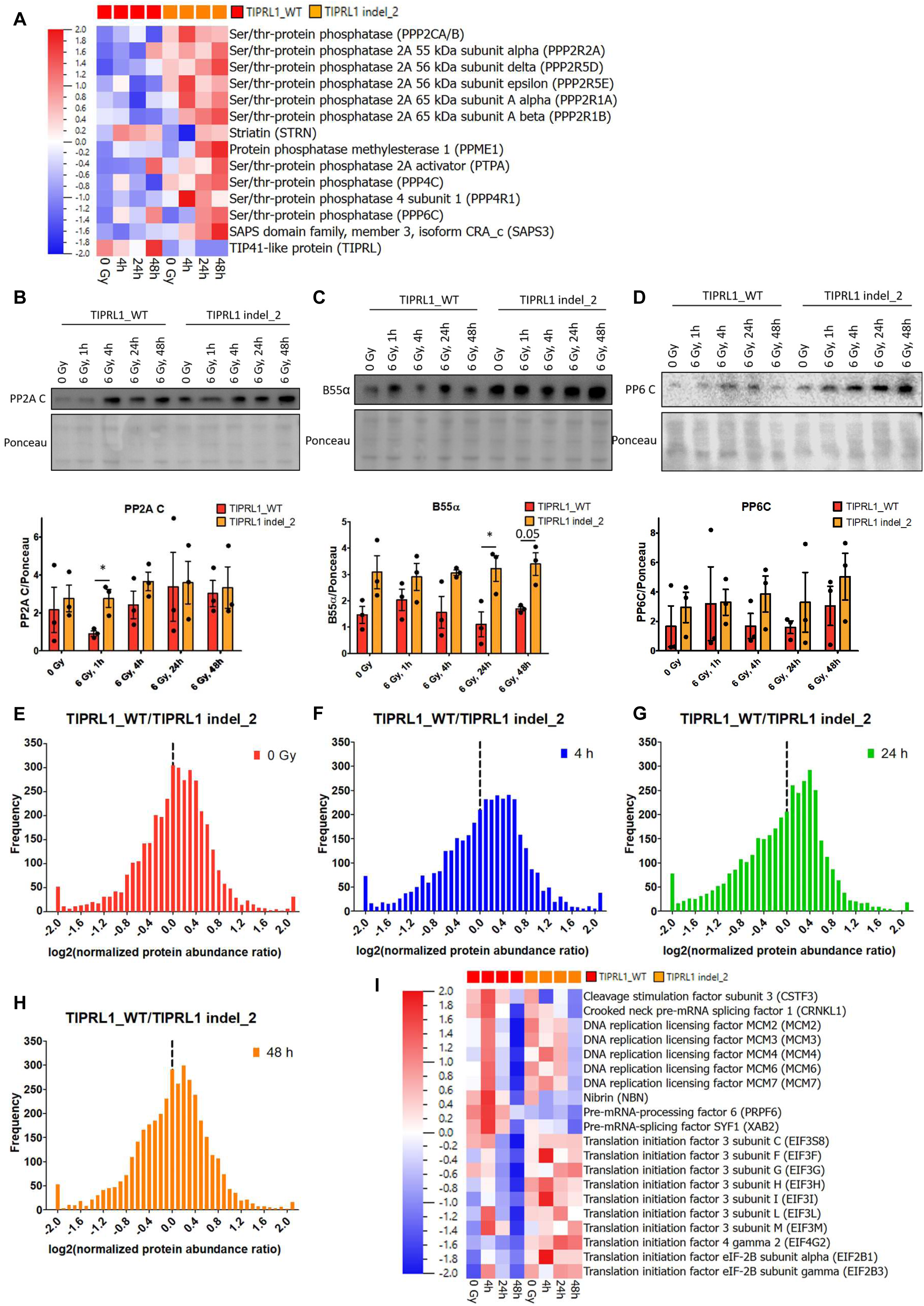
Effect of TIPRL1 depletion on the global SQD9 proteome in basal and irradiated conditions. Unbiased proteomics analysis was performed on total lysates from SQD9 cells with TIPRL1 WT expression or *TIPRL* indel mutation without irradiation or 4h, 24h and 48h after exposure to 6 Gy irradiation (n=1). The total amount of proteins loaded on LC-MS/MS in each condition was very similar, displaying a CV (variation coefficient) of 12% for the average, and of 19% for the standard deviation of the raw protein abundance between all 8 samples. **A** Heat map showing normalized protein abundances of all identified PP2A-like phosphatase-related proteins in the indicated cell lines and conditions. **B – D** Validation of expression of PP2A C (**B**), B55α (**C**) and PP6C (**D**) in TIPRL1_WT cells vs TIPRL1 indel_2 cells. Western blots of total lysates without irradiation, or 1h, 4h, 24h or 48h after exposure to 6 Gy were developed using a primary PP2A C, B55α and PP6C antibody. Ponceau was used as loading control. A representative blot is shown (upper panel). Quantifications represent mean ± SEM of n=3 (t-test, *: p<0.05) (lower panel). **E – H** Frequency distribution of the logarithms of normalized protein abundance ratios between TIPRL1_WT and TIPRL1 indel_2 cell lysates without (**E**) or 4h (**F**), 24h (**G**) and 48h (**H**) after exposure to 6 Gy irradiation. **I** Heat map showing normalized protein abundances of enriched genes (David functional annotation clustering analysis) after rank regression (p<0.05) with a negative slope of the regression line of normalized protein abundances obtained from TIPRL1_WT cells (4h, 24h and 48h post-RT).

To obtain an overall view of potential protein expression changes in function of irradiation and how this might be affected by loss of TIPRL1, we drew a frequency distribution of the logarithms of normalized protein abundance ratios between TIPRL1_WT and TIPRL1 indel_2 cell lysates at each time point (**Fig 4E-4H**). Clearly, the obtained distributions were slightly skewed to values above 0 in basal conditions (0 Gy) (**Fig 4E**) and even more obviously at 4h and 24h post-RT (**Fig 4F, 4G**), while at 48h post-RT the distribution was rather skewed to values below 0 (**Fig 4H**). These expression kinetics might be explained by (1) proteins whose expression is initially increased upon irradiation, specifically in the WT cells (4h and at 24h post-RT), and is subsequently decreased again by 48h post- RT to levels that are even lower than in basal conditions – and/or (2) proteins whose expression is initially decreased upon irradiation, specifically in TIPRL1 depleted cells (4h and 24h post-RT), and is subsequently increased by 48h post-RT to levels that are even higher than in basal conditions. In order to identify which proteins were actually following this expression kinetics, we subjected normalized protein abundances obtained from TIPRL1_WT cells (4h, 24h and 48h post-RT) to rank regression (p<0.05) with a negative slope of the regression line as a requirement, while normalized protein abundances obtained from TIPRL1 indel_2 cells (4h, 24h and 48h post-RT) were subjected to rank regression (p<0.05) with a positive slope of the regression line. We identified 300 protein entries fulfilling this criterion for TIPRL1_WT cells, and 235 protein entries for TIPRL1 indel_2 cells (**Appendix Table S2**). The obtained protein lists were subsequently subjected to David functional annotation clustering analysis using the total experimental dataset (3793 entries) as a background. For TIPRL1_WT cells (negative slope of the regression line), 300 protein entries could be linked to 256 David gene IDs resulting in only 4 high-stringency clusters with enrichment scores >2.0, corresponding to 20 genes (**Fig 4I**) involved in ‘DNA replication and repair’, ‘translation initiation’ and ‘mRNA processing’. For TIPRL1 indel_2 cells (positive slope of the regression line), 235 protein entries could be linked to 195 David gene IDs and 4 high-stringency clusters, corresponding to 79 genes that showed no relationship to replication, transcription or translation processes, but were related to ‘cell-adhesion’, ‘small GTPases’ and ‘histone H2A’ (**Appendix Fig S2**). Moreover, this heat map (**Appendix Fig S2**) but not the former one (**Fig 4I**) indicated that normalized protein abundances at 0 Gy were systematically higher in TIPRL1_WT compared to TIPRL1 indel_2 cells, potentially explaining the skewed shape of the frequency distribution graph in non-irradiated cells (**Fig 4E**). In summary, both groups of proteins abiding by the altered expression kinetics in irradiated TIPRL1_WT or TIPRL1 indel_2 cells were clearly linked to different cellular processes. The former group (‘DNA replication and repair’, ‘translation initiation’ and ‘mRNA processing’) was specifically affected upon irradiation in TIPRL1-depleted cells, while the latter group (‘cell adhesion’, ‘small GTPases’ and ‘histone H2A’) was already affected under basal conditions and was not further altered upon irradiation. Finally, when performing a rank regression analysis of TIPRL1_WT and TIPRL1 indel_2 proteins with slope requirements that were opposite to those of the expected expression kinetics after irradiation, David analysis of these protein lists did not yield any high-stringency cluster with enrichment score >2.0, further underscoring the relevance of the above described enriched clusters.

### TIPRL1 shows an increased interaction with nucleosomal histones and DDR regulators DNA-PKcs and RAD51 upon radiation, while interaction with PP2A, PP4 and PP6 is not modulated

To identify putative alterations in the TIPRL1 interactome upon irradiation, we performed AP-MS with 3xFLAG-tagged TIPRL1 from Rescue TIPRL1_WT cells as the bait. TIPRL1_WT cells lacking the expression of any tagged proteins were used as negative control. FLAG pulldowns were executed on lysates from irradiated and non-irradiated cells at 1h and 4h post-RT (6 Gy). Given conflicting data on the effects of phosphatase inhibitors (PPIs), such as okadaic acid, on the interaction between TIPRL1 and PP2A (Smetana & Zanchin, 2007; Wu et al, 2017), we prepared cell lysates and performed anti-FLAG IPs in the presence as well as in the absence of PPIs.

Overall, 140 cellular TIPRL1 interactors were identified in at least one of these conditions following our stringent MS-related selection criteria (*cfr.* Materials and Methods) (**Appendix Table S3**). Intriguingly, we identified significantly more TIPRL1 interaction partners in the presence of PPIs than in the absence of PPIs in a given condition (χ^2^ test: all p-values <0.05) (**Fig 5A, Appendix Table S3**). This finding exceeded our expectations considering the anticipated effects of PPIs on the interaction with PP2A alone. Likewise, the number of interactors in irradiated cells showed a significant decrease in the presence of PPIs at 4h post-RT, while the opposite was seen in the absence of PPIs at 1h and 4h post-RT (**Fig 5A, Appendix Table S3**). These results suggest phosphorylation-dependent changes in the TIPRL1 interactome.

**Figure 5.**
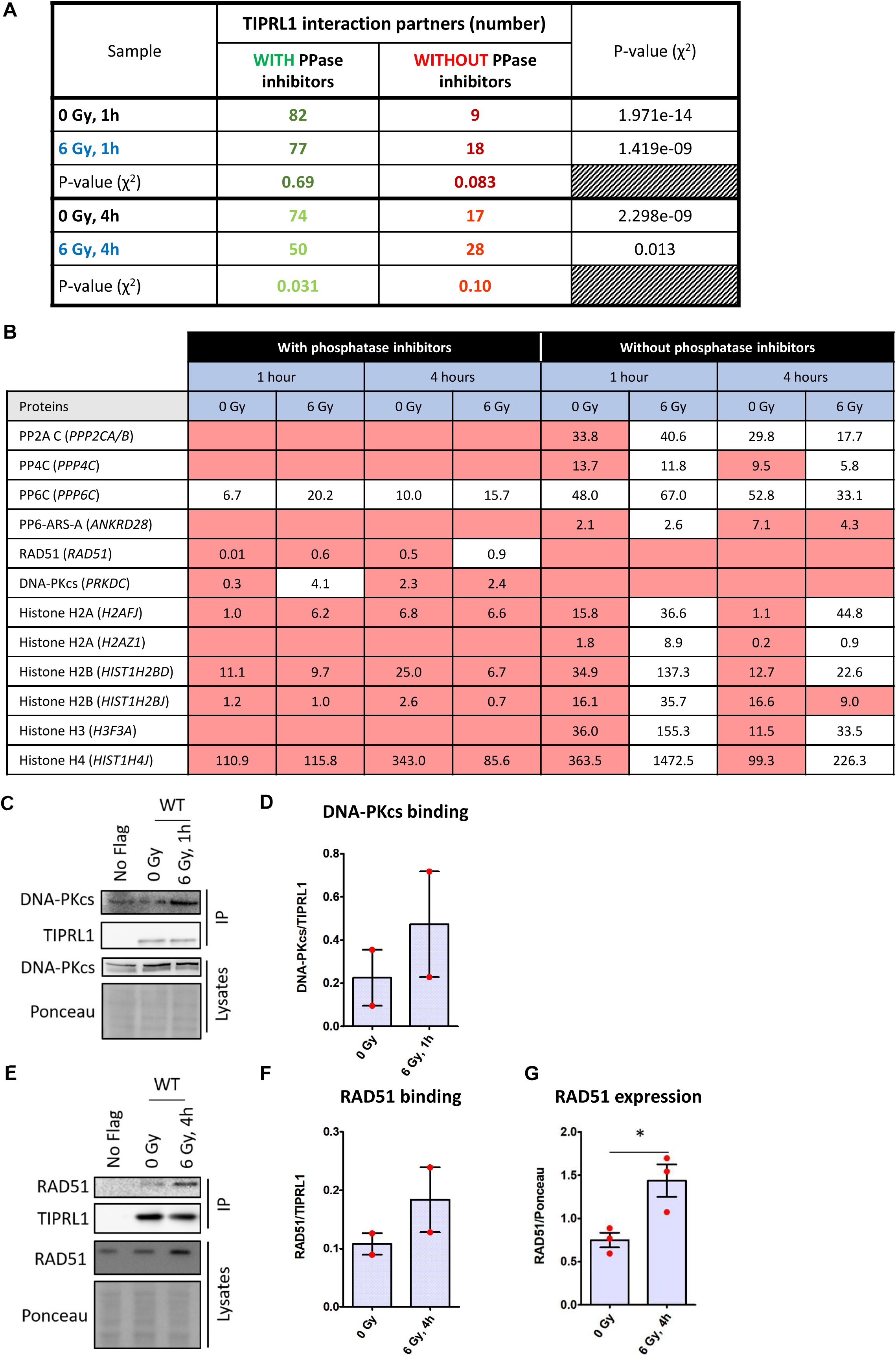
TIPRL1 WT interactome upon irradiation. **A** TIPRL1 interactome was determined in the absence and in the presence of phosphatase inhibitors (PPIs) by AP-MS. Chi-square test was performed to assess overall differences in detected TIPRL1 interaction partners with or without PPIs, and with or without irradiation for 1h or 4h after 6 Gy. **B** Select TIPRL1 interaction partners are presented with their corresponding protein abundance normalized to the protein abundance of immunoprecipitated TIPRL1. Cells marked in red are considered no TIPRL1 interactors based on stringent MS-related criteria (c*fr* Materials and Methods for more details). Empty cells indicate that the protein was not detected in the corresponding condition. **C, D** Validation of increased DNA-PKcs binding to TIPRL1 upon irradiation. Protein lysates of non-irradiated (0 Gy) or irradiated (6 Gy, 1h post-RT) TIPRL1_WT (No Flag), and Rescue TIPRL1_WT cells were prepared, and subjected to anti-FLAG immunoprecipitation (IP). Representative blot of DNA-PKcs expression and binding to TIPRL1 as a function of irradiation (**C**) with corresponding quantification of TIPRL1–DNA-PKcs binding (n=2) (**D**). Ponceau was used as loading control. **E – G** Validation of increased RAD51 binding to TIPRL1 upon RT. Protein lysates of non-irradiated (0 Gy) or irradiated (6 Gy, 4h post-RT) TIPRL1_WT (No Flag), and Rescue TIPRL1_WT cells were prepared, and subjected to anti-FLAG immunoprecipitation (IP). Representative blot of RAD51 expression and binding to TIPRL1 as a function of irradiation (**E**), with corresponding quantification of TIPRL1–RAD51 binding (n=2) (**F**) and RAD51 expression (n=3) (**G**). Significant differences (mean ± SEM) are indicated with *: p<0.05 (t-test).

TIPRL1 interaction with the catalytic subunits of all three PP2A-like phosphatases (PP2A C, PP4C and PP6C) was detected in the absence of PPIs (**Fig 5B**), while in the presence of PPIs, surprisingly, only the interaction with PP6C was still detected (**Fig 5B**). No additional PP2A-like phosphatase subunits were found in the interactome other than PP6-ARS-A (*ANKRD28*), which appeared as an interactor at 1h post-RT in the absence of PPIs (**Fig 5B**). No major RT-dependent modulation of PP2A C, PP4C or PP6C binding to TIPRL1 could be established in any of the tested conditions (**Appendix Table S4**). In contrast, some novel TIPRL1 interactors with reported functions in the DDR showed RT-dependent or RT-modulated binding to TIPRL1 (**Appendix Table S4**), which made them promising candidates for further follow up.

Specifically, RAD51, a DNA repair protein involved in homologous recombination (HR) (Baumann & West, 1998; Lee & Paull, 2021), and the DNA damage sensor kinase DNA-PKcs (Blackford & Jackson, 2017) were found to be novel TIPRL1 interaction partners in the presence of PPIs (**Fig 5B; Appendix Table S3, Table S4**). These interactions and their increase following RT were independently confirmed by immunoblotting for endogenous DNA-PKcs (**Fig 5C, 5D**) and RAD51 (**Fig 5E, 5F**) following anti-FLAG TIPRL1 immunoprecipitation from irradiated Rescue TIPRL1_WT cells. However, both MS and immunoblot data indicated a more reproducible interaction for RAD51 in comparison to DNA-PKcs. Notably, for RAD51, total protein levels also increased in irradiated cells (**Fig 5E, 5G**), potentially explaining why an increased interaction with TIPRL1 was observed. However, despite this increased interaction with TIPRL1, neither the irradiation-induced modulation of RAD51 Thr309 phosphorylation nor that of DNA-PKCs Ser2056 phosphorylation was affected by the loss of TIPRL1 (**Fig EV2**).

Another group of novel TIPRL1 interactors of potential interest were the nucleosomal histones H2A, H2B, H3 and H4. They were all independently detected as TIPRL1 interactors upon RT and in the absence of PPIs (**Fig 5B, Appendix Table S3, Table S4**) - suggesting they are genuine TIPRL1 interactors in irradiated cells and their binding may be modulated by a putative phosphorylation.

### ATM kinase phosphorylates TIPRL1 on Ser265 upon RT

Given the observed differences in the TIPRL1 interactome in the presence as opposed to the absence of PPIs, we hypothesized that TIPRL1 itself might become phosphorylated upon irradiation. We performed unbiased MS-based analysis of 3xFLAG-tagged TIPRL1 isolated from irradiated cells (at 1h post-RT) and identified two tryptic monophosphorylated peptides encompassing the same region of TIPRL1: 258-IDPNPADSQK-267 and 258-IDPNPADSQKSTQVE-272 (**Fig 6A**). According to the Proteome Discoverer PhosphoRS node, the phosphorylation resided exclusively at Ser265. To further examine the kinetics of Ser265 phosphorylation, we performed a targeted MS-based phospho analysis for Ser265 encompassing peptides on 3xFLAG-tagged TIPRL1 isolated from irradiated cells at different time points after irradiation. As a result, maximal TIPRL1 Ser265 phosphorylation was retrieved at 2h post-RT (**Fig 6B**). To validate Ser265 as the main TIPRL1 phosphorylation site, we introduced a non- phosphomimic TIPRL1 S265A mutant or a presumed phosphomimic S265D mutant in the TIPRL1 indel_1 cells and confirmed the expression of the mutants by immunoblotting (**Fig 6C, 6D**). When applying the same targeted MS-based approach on TIPRL1 isolated from all three rescue cell lines, we only observed phosphorylation in the WT Ser265 encompassing peptide (**Fig 6E**). Importantly, Ser265 phosphorylation of WT TIPRL1 could only be detected in the presence of PPIs: when PPIs were omitted from the assays, Ser265 phosphopeptides could no longer be identified in the targeted MS analyses.

**Figure 6.**
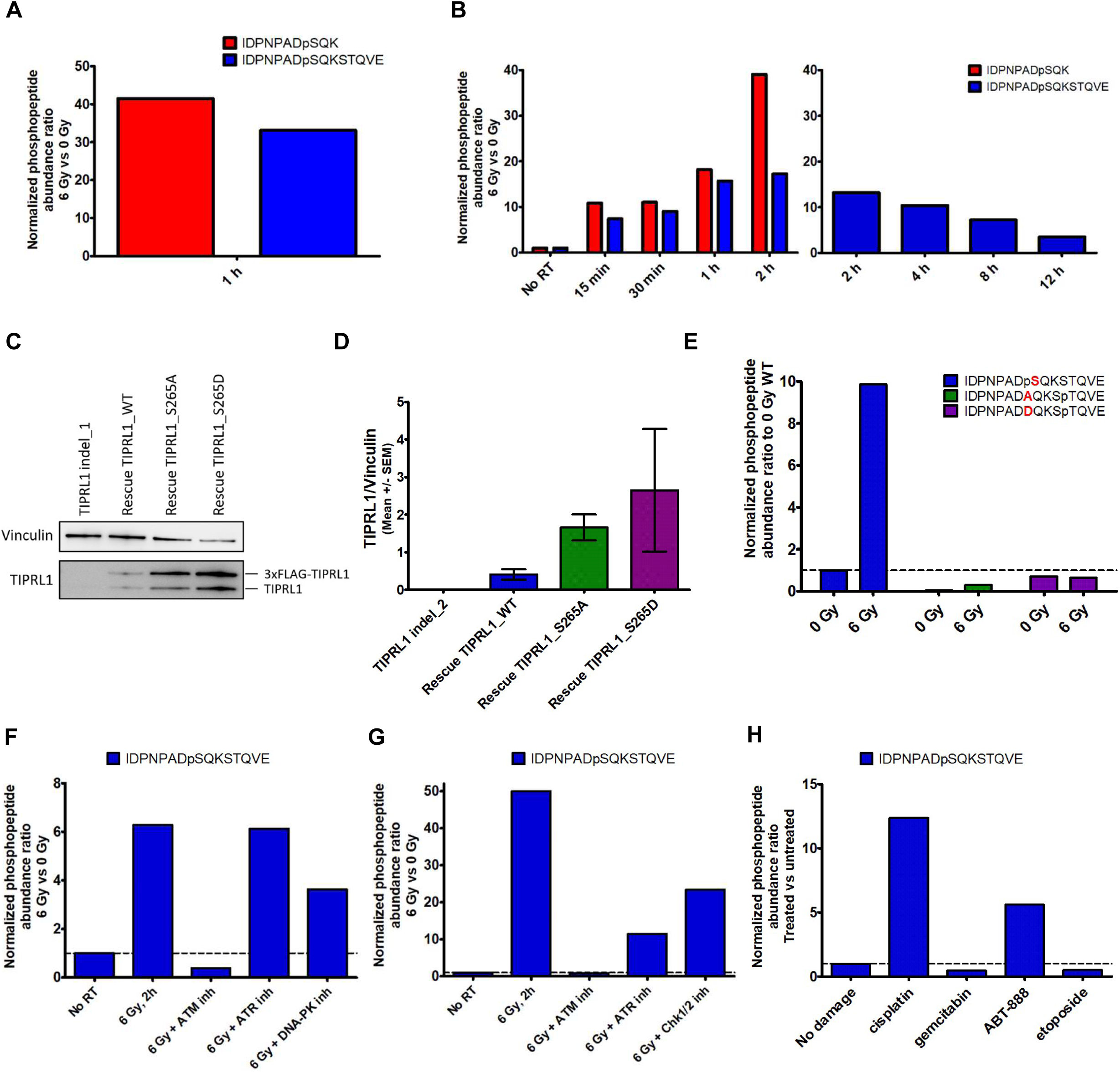
DNA damage induces TIPRL1 phosphorylation at Ser265 by ATM kinase. **A** Unbiased phosphoproteomics analysis of 3xFLAG-tagged TIPRL1 immunoprecipitates from Rescue TIPRL1_WT SQD9 cells without and with 1h of 6 Gy irradiation (n=1). **B** Targeted MS analysis of TIPRL1 Ser265 phosphorylation as a function of time after irradiation (n=1). **C, D** Transduction of TIPRL1 indel_1 cells with a FLAG-tagged TIPRL1 phosphomimic (S265D) or non- phosphomutant (S265A) expressing lentivirus. As the start codon of the mutant TIPRL1 coding sequence was not removed, some untagged mutant TIPRL1 is also re-expressed. Representative blot of the successful viral transduction (**C**) and the corresponding quantification of TIPRL1 expression (FLAG-tagged plus untagged) normalized to Vinculin (n=3, mean ± SEM, one-way ANOVA followed by Tukey’s multiple comparison test) (**D**). **E** Targeted MS analysis of WT versus S265A and S265D TIPRL1 phosphorylation as a function of irradiation (2h post 6 Gy) (n=1). **F, G** Targeted MS analysis of TIPRL1 Ser265 phosphorylation after pretreatment of Rescue TIPRL1_WT SQD9 cells for 2h with 1.5 µM AZD1390 (ATM inh), 0.5 µM AZD6738 (ATR inh), 0.5 µM AZD7648 (DNA-PK inh) or 0.25 µM AZD7762 (Chk1/2 inh), followed by exposure to 6 Gy irradiation and collection (after 2h). Two independent experiments are shown. **H** Targeted MS analysis of TIPRL1 Ser265 phosphorylation after treatment of Rescue TIPRL1_WT SQD9 cells for 2h with DMSO, 0.5 µM cisplatin, 5 nM gemcitabine, 20 µM ABT-888 or 5 µM etoposide (n=1).

As Ser265 resides in an ATM/ATR/DNA-PK kinase consensus sequence (pS/TQ-site) (Kim et al, 1999; O’Neill et al, 2000), we next pretreated Rescue TIPRL1_WT SQD9 cells for 2h with ATM (AZD1390), ATR (AZD6738), or DNA-PK (AZD7648) inhibitors to identify the TIPRL1 kinase. Only in the presence of the ATM inhibitor, TIPRL1 phosphorylation was completely abolished at 2h post-RT (**Fig 6F**). To further exclude that the observed decrease in TIPRL1 phosphorylation was the result of the inhibition of a downstream ATM kinase, we also pretreated irradiated Rescue TIPRL1_WT cells with the Chk1/Chk2 kinase inhibitor AZD7762, but could not demonstrate strong modulation of TIPRL1 phosphorylation (**Fig 6G**). We concluded that ATM directly phosphorylates TIPRL1 at Ser265 in irradiated SQD9 cells.

To further establish whether TIPRL1 phosphorylation was exclusively triggered by irradiation-induced DNA damage, we also assessed the effects produced by the addition of other DNA damaging agents (**Fig 6H**). Surprisingly, we found some specificity, as cisplatin and the PARP inhibitor ABT-888 both induced TIPRL1 Ser265 phosphorylation, while gemcitabine and etoposide did not (**Fig 6H**).

### TIPRL1 requires RT-induced Ser265 phosphorylation to mediate radiotherapy resistance

To elucidate the importance of the ATM-mediated TIPRL1 Ser265 phosphorylation in the RT response in SQD9 cells, 2D colony assays were performed with TIPRL1 indel_1, Rescue TIPRL1_S265A and Rescue TIPRL1_S265D cells. While all three cell lines showed similar plating efficiencies (**Fig 7A**), TIPRL1 S265D mutant expressing cells showed significantly increased radiotherapy resistance as opposed to TIPRL1 indel_1 and TIPRL1 S265A mutant expressing cells (**Fig 7B**), further underscored by increased dose enhancement factors (**Fig 7C**). In addition, Rescue TIPRL1_S265D cells (**Fig 7C**) showed a similar DEF as Rescue TIPRL1_WT cells (**Fig 2E**) relative to TIPRL1 indel_1 (2.0 vs 1.9 at 6 Gy). Consistently, the radioresistant Rescue TIPRL1_S265D cell line retained fewer micronucleated cells at 48h post-RT (6 Gy) than the more radiosensitive cell lines lacking TIPRL1 expression or expressing the non-phosphorylatable TIPRL1 S265A mutant (**Fig 7D**). Thus, TIPRL1 requires Ser265 phosphorylation by ATM to mediate RT resistance in HNSCC cells.

**Figure 7.**
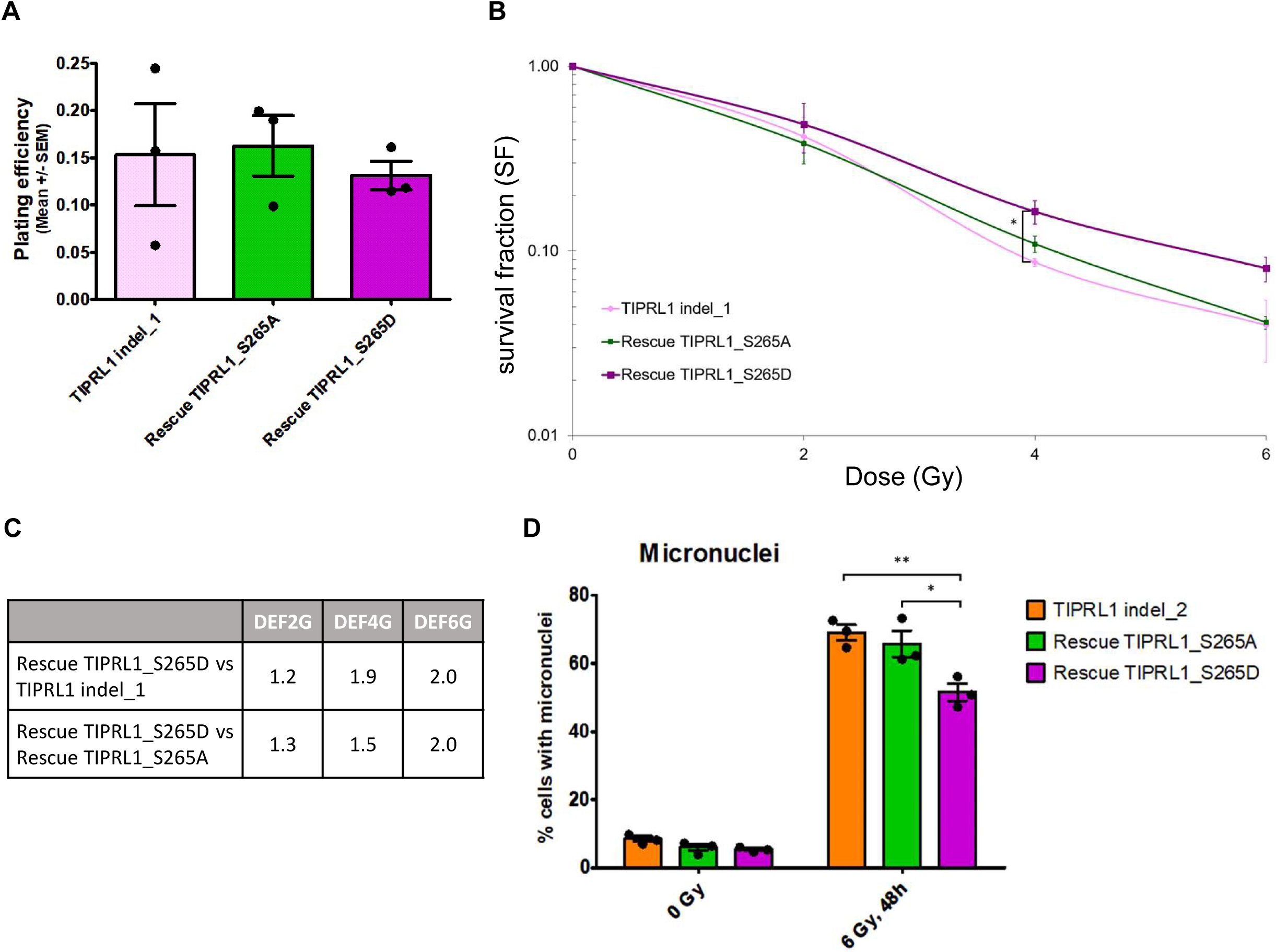
Constitutive phosphorylation-mimicking TIPRL1 mutant (S265D) induces radiotherapy resistance. **A** Calculation of plating efficiency as the ratio between the number of colonies and the number of cells seeded (n=3, mean ± SEM, one-way ANOVA followed by Tukey’s multiple comparison test) for TIPRL1 indel_1, Rescue TIPRL1_S265A and Rescue TIPRL1_S265D cells. **B** Survival fractions (logarithmic scale) of TIPRL1 indel_1, Rescue TIPRL1_S265A and Rescue TIPRL1_S265D cells after different doses of RT (0, 2, 4 and 6 Gy) shown as the mean clonogenic survival ± SEM (n=3, one-way ANOVA followed by Tukey’s multiple comparison test, *: p<0.05). **C** Dose Enhancement factors (DEF) for different doses of RT (2, 4 and 6 Gy) expressed as the ratios of the survival fractions of cells expressing the phosphomimic (S265D) *vs* either TIPRL1 indel_1 cells or *vs* cells expressing the non-phosphomutant (S265A). **D** Assessment of the percentage of micronucleated cells from TIPRL1 indel_2 SQD9 cells or those expressing the non-phosphomutant (S265A) or phosphomimic (S265D) after 6 Gy irradiation. DAPI staining was performed 48h after irradiation (n=3, mean ± SEM, one-way ANOVA followed by Tukey’s multiple comparison test, *: p<0.05, **: p<0.01).

As our TIPRL1 interactomics data demonstrated RT-dependent modulation of TIPRL1 binding to RAD51, DNA-PKCs and histones, but not to PP2A-like phosphatases (**Fig 5B-5G**), we next asked whether this modulation was dependent on TIPRL1 Ser265 phosphorylation. Western blot analysis showed that PP2A C (**Fig EV3A**) and PP6C (**Fig EV3B**) still bound equally well to TIPRL1 regardless of Ser265 mutation or RT-induced DNA-damage, confirming the lack of modulation of their binding by RT and by Ser265 phosphorylation. PP4C could not be validated due to lack of suitable antibodies. On the other hand, while DNA-PKcs (**Fig EV3C**) as well as RAD51 binding (**Fig EV3D**) to TIPRL1 S265A did no longer increase upon RT, a similar result was obtained with TIPRL1 S265D (**Fig EV3C, EV3D**). As TIPRL1 S265D behaved as a true phospho-mimic in our phenotypic assays (**Fig 7B**, **7D**), we conclude that increased TIPRL1 binding to RAD51 or DNA-PKcs is not causally involved in the increased radioresistance of Rescue TIPRL1_S265D cells, and, perhaps, may require the full integrity of Ser265. To further confirm and extend these observations, we repeated the unbiased AP-MS interactomics analysis with 3xFLAG-tagged TIPRL1 S265A and S265D mutants. Neither RAD51, nor DNA-PKcs could be identified in the TIPRL1 S265A and S265D interactomes, irrespective of irradiation, while all PP2A- like phosphatases were still found to bind both TIPRL1 mutants (**Appendix Table S3, Table S4**) - confirming and extending our targeted immunoblot data (**Fig EV3**). Notably, binding of all four nucleosomal histones H2A, H2B, H3 and H4 to the TIPRL1 S265A mutant was significantly induced upon irradiation in the absence as well as in the presence of PPIs. Conversely, the binding of all nucleosomal histones to TIPRL1 S265D showed the opposite trend (**Fig 8A, 8B, Appendix Table S3, Table S4**). Since increased histone–TIPRL1 WT binding was exclusively seen in irradiated cells in the absence of PPIs (not preserving Ser265 phosphorylation), these findings would not only suggest a clear binding preference of non-phosphorylated TIPRL1 to the histones in irradiated cells, but also an additional facilitation of the binding by a RT-induced factor, such as *e.g*. a RT-induced modification of the histones.

**Figure 8.**
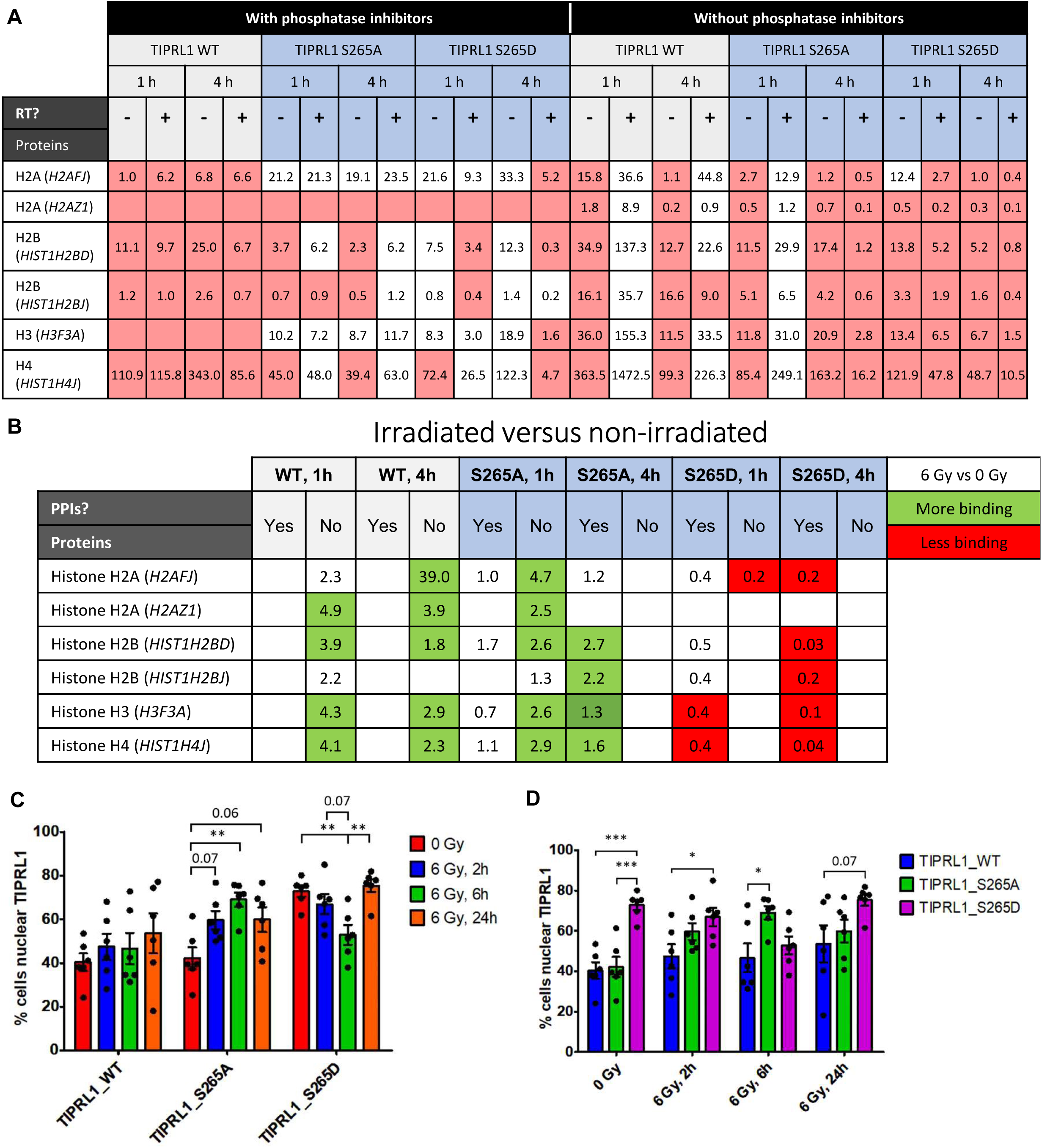
Effect of irradiation-induced TIPRL1 phosphorylation on nucleosomal histone binding and nuclear/cytoplasmic TIPRL1 distribution. **A** TIPRL1 WT, TIPRL1 S265A and TIPRL1 S265D interactomes were determined by AP-MS, in the absence and in the presence of phosphatase inhibitors (PPIs), and in the indicated conditions (RT: 6 Gy). A selection of TIPRL1 interaction partners, i.e. all nucleosomal histones, is presented with their corresponding protein abundance normalized to the protein abundance of immunoprecipitated TIPRL1. Cells marked in red are considered no TIPRL1 interactors based on stringent MS-related criteria (c*fr* Materials and Methods for more details). Empty cells indicate that the protein was not detected in the corresponding condition. **B** Overview of the effects of irradiation on TIPRL1-histone interactions. Normalized abundance ratios were calculated for all histones in each condition after treatment with 6 Gy versus 0 Gy. Conditions marked in green represent increased TIPRL1 binding upon irradiation, conditions marked in red indicate decreased binding after RT. **C, D** SQD9 cells expressing 3xFLAG-tagged WT TIPRL1, TIPRL1 S265A mutant, or TIPRL1 S265D mutant were exposed to 6 Gy irradiation and fixed for immunofluorescence staining at different time points after radiotherapy with the FLAG-tag antibody to determine the percentage of cells with nuclear TIPRL1. Significant differences are indicated with *: p<0.05, **: p<0.01, ***: p<0.001 (n=1, 6 technical replicates, mean ± SEM, one-way ANOVA followed by Tukey’s multiple comparison test).

Additional immunofluorescence analysis of 3xFLAG-tagged TIPRL1 subcellular localization performed in Rescue TIPRL1_WT, Rescue TIPRL1_S265A and Rescue TIPRL1_S265D cells indicated an increased presence of the S265D mutant in the nucleus under basal conditions (**Fig 8C, 8D**). Upon RT (6 Gy), the fraction of cells with nuclear presence of S265D TIPRL1 decreased, and returned to basal levels at 24 hours post-RT. The opposite trend was seen for S265A TIPRL1, showing an increased nuclear presence at 2h and 6h post-RT, and declining again at 24h. For TIPRL1 WT expressing cells, the fraction of cells with nuclear presence of WT TIPRL1 was not modulated (**Fig 8C, 8D**). These data would suggest a balance of TIPRL1 WT localization between the cytoplasm and the nucleus regardless of irradiation, which might become (artificially) distorted upon Ser265 substitution in irradiated conditions - and that might be related to the observed increased binding of TIPRL1 S265A to the nucleosomal histones (**Fig 8B**) and the observed decreased histone binding of TIPRL1 S265D (**Fig 8B**) in irradiated cells.

## Discussion

Understanding the mechanistic foundation of tumor cell radioresistance is of vital importance to enhance treatment outcomes and prevent tumor recurrence in patients with HNSCC. Activating cell cycle checkpoints to trigger the DDR and promote DNA repair is one of the major mechanisms of tumor cell survival upon exposure to RT.

In the present study, we exposed a role for TIPRL1, a functionally poorly understood binding partner and inhibitor of PP2A-like phosphatases, in the RT response of HNSCC cells. Upon analysis of two independent patient cohorts, we showed that (1) on average, TIPRL1 expression is significantly increased in HNSCC tissue as opposed to non-tumor tissue, and (2) within tumors, high mRNA expression or immunostaining of TIPRL1 correlated with decreased locoregional control and lower survival, specifically for patients treated with radiotherapy. Although the small number of samples in our immunohistochemical analysis certainly represents a limitation, this still suggests that high TIPRL1 expression may be predictive for poorer outcomes of RT. Whether increased TIPRL1 expression is relevant in other cancers remains to be determined, although three recent studies in pancreatic and liver cancer also linked increased TIPRL1 expression to decreased patient survival (Jun et al, 2019, 2021; van Pelt et al, 2023). Together with our own observations in HNSCC, this would add TIPRL1 to the growing list of clinically relevant PP2A inhibitors with a prognostic value in cancer (Khanna & Pimanda, 2016; Meeusen & Janssens, 2018), and specifically in HNSCC, identify it as a predictor of RT resistance.

Importantly, we underscored these clinical findings in our cellular studies, demonstrating substantial radiosensitization of SQD9 cells upon TIPRL1 depletion. This result is in contrast with a study in Hela cells (Rosales et al, 2015), but aligns well with reports in *Saccharomyces cerevisiae* (Gingras et al, 2005) and *Candida albicans* (Feng et al, 2016), in which deletion of ScTip41 or CaTip41 sensitized to cisplatin. In the latter two studies cisplatin sensitization was also observed upon deletion of Pph3 (orthologue of human PP4C), which seems at odds with Tip41’s alleged function as a Pph3 inhibitor. Although it remains to be determined whether the role of TIPRL1 in the DDR in our HNSCC model involves its PP2A-like phosphatase modulatory ability, our work demonstrated a clear dependence on its DNA damage-induced phosphorylation by ATM kinase.

McConnell et al. (2007) were the first to imply TIPRL1 in ATM-dependent signaling in mammalian cells: upon siRNA-mediated knockdown of TIPRL1 in HEK293 cells, they observed downregulation of a 32-kDa-protein that was recognized by ATM/ATR phosphosite motif antibodies (McConnell et al, 2007). Based on our current data, we would argue that this protein may very likely have been TIPRL1 itself. We indeed identified TIPRL1 as a novel substrate of ATM in irradiated SQD9 cells, with maximal phosphorylation observed around 2h post-RT, and with Ser265, residing in an ATM consensus motif (pSQ), as main phosphorylation site. We also found Ser265 phosphorylation upon treatment of the cells with cisplatin and a PARP inhibitor, but not upon addition of gemcitabine or etoposide. As all these treatments in principle should activate ATM, this may reflect additional regulation of TIPRL1- directed ATM activity depending on the type of DNA lesion triggered. Although Ser265 phosphorylation had been identified before in large-scale screening studies (Boeing et al, 2016; Matsuoka et al, 2007; Sharma et al, 2014; Zhou et al, 2013), its specific induction upon RT, cisplatin and PARP inhibitor exposure is a novel observation. We cannot entirely exclude that Thr269 might not represent an additional, albeit very marginal, ATM phosphorylation site as well. Although the Proteome Discoverer PhosphoRS node attributed phosphorylation of the monophosphorylated 258- IDPNPAD**S**QKS**T**QVE-272 peptide for 100% to the Ser265 residue, manual interpretation of the MS/MS information obtained from Skyline could not completely rule out phosphorylation at Thr269: based on the summed intensities of MS/MS fragments exclusive for either the Ser265 or Thr269 phosphorylated species, the contribution of Thr269 phosphorylation would be <20%. However, since (1) the 258-IDPNPAD**<u>S</u>**<u>Q</u>KS**<u>T</u>**<u>Q</u>VE-272 peptide was never identified as a double phosphorylated species, (2) phosphorylation of the Ser265 phospho ablative or phosphomimic peptides (258- IDPNPAD**<u>(A/D)</u>**QKS**T**QVE-272) isolated from irradiated TIPRL1 S265A or D mutant expressing cells was always lower than basal (0 Gy) phosphorylation of WT peptides, and (3) the Thr269-encompassing ATM consensus site is not conserved in mouse (TP instead of TQ), we considered Thr269 phosphorylation upon RT as highly unlikely.

According to crystallographic studies, Ser265 resides in an unstructured domain nearby the C- terminus of the 272-amino-acid TIPRL1 protein, a region that is not involved in PP2A C binding nor in modulation of PP2A inhibition (Scorsato et al, 2016; Wu et al, 2017). Consistently, we found that TIPRL1 binding to PP2A, PP4 or PP6 was not modulated by RT in our assays (although binding could only be shown in the absence of PPIs for PP2A and PP4), and that PP2A, PP4 and PP6 still bound equally well to TIPRL1 S265A and S265D mutants as to WT TIPRL1, irrespective of irradiation. Likewise, catalytic inhibition of a purified PP2A dimer by (unphosphorylated) recombinant TIPRL1 was not affected by TIPRL1 S265D substitution in an *in vitro* phosphatase assay (**Appendix Fig S3**). Overall, these findings imply that ATM-mediated Ser265 phosphorylation of TIPRL1 does not affect its binding to the PP2A-like phosphatases, nor its inhibitory potential toward PP2AD. As a bonus result, our interactomics data also suggested for the first time PP2A-like phosphatase-specific changes in TIPRL1-binding mode and thus, potentially, in TIPRL1-dependent modulation of PP2A-like phosphatase activity - as PP2A-TIPRL1 and PP4-TIPRL1 binding did not occur in the presence of PPIs, while PP6-TIPRL1 binding remained unaffected. In addition, in accordance with previous work (Gingras et al, 2005), the catalytic subunits of PP2A, PP4 and PP6 were the main phosphatase constituents present in the TIPRL1 interactome regardless of irradiation. No other phosphatase regulatory subunits (except PP6 ANKRD28, 1h post-RT in the absence of PPIs), nor alpha4, could consistently be identified. On the other hand, the unbiased proteomics analysis of TIPRL1_WT and TIPRL1 indel_2 cells identified increased expression of all detected phosphatase subunits/regulators in TIPRL1-depleted SQD9 cells, which seems at odds with its proposed phosphatase stabilizing role in ‘stressed’ cells, and is opposite to the described role of alpha4 under stress conditions (Kong et al, 2009). So, exactly how TIPRL1 regulates PP2A, PP4 and PP6 within cells still remains insufficiently clear.

Our proteomics data further provided insights into the role of TIPRL1 in TOR signaling and the initiation of translation. Although originally identified as a negative regulator of TOR signaling in yeast (Jacinto et al, 2001), contrasting results were reported in mammalian cells (Nakashima et al, 2013). We found several subunits of the eIF2, eIF3 and eIF4 complexes significantly enriched in TIPRL1_WT cells within the set of proteins showing a negative rank regression in irradiated TIPRL1_WT cells, leading to our current speculation that (phosphorylated?) TIPRL1 might actually promote translation initiation of specific DNA repair proteins (Mei et al, 2022; Tamaddondoust et al, 2022). Likewise, the RT-induced expression modulation of several proteins involved in mRNA processing and alternative splicing was no longer found in TIPRL1-depleted cells. As part of the DDR also involves changes in pre-mRNA alternative splicing of specific gene products needed for checkpoint activation or DNA repair (Chen et al, 2017; Lee & Paull, 2021), this could further sustain a positive role for TIPRL1 in the DDR. Another significantly enriched cluster in irradiated TIPRL1_WT cells encompassed several subunits of the Mini Chromosome Maintenance (MCM2-7) complex, a DNA helicase involved in replication fork assembly and DNA replication. In case of DNA damage and upon S-phase checkpoint activation, fork progression is stopped by promoting stable MCM2-7 binding to the fork during DNA repair (Li & Xu, 2019). A last protein that was specifically enriched in irradiated TIPRL1_WT cells was nibrin, also known as Nijmegen Breakage Syndrome 1 (NBS1), a DNA repair protein of the MRN complex that is involved in ATM activation and subsequent HR repair (Lee & Paull, 2021). All these findings, together with our observations that TIPRL1-depleted cells showed a longer maintained checkpoint activation and displayed more micronuclei formation at 48h post-RT, underscore a positive role for TIPRL1 in the DDR, particularly by promoting DNA repair processes.

This general hypothesis was further corroborated by our interactomics studies, which identified several novel TIPRL1 interactors involved in DNA repair, and for which the interaction with TIPRL1 was modulated by irradiation. One of these proteins was DNA-PKcs, a major driver of Non- Homologous End Joining (NHEJ) repair of dsDNA breaks (DSBs) (Blackford & Jackson, 2017). TIPRL1– DNA-PKcs binding, albeit weak, was no longer detectable by Ala or Asp substitution of TIPRL1 Ser265 in our AP/MS assays, and barely by immunoblotting. For TIPRL1–RAD51 binding, similar data were found, although RAD51 interaction with TIPRL1 S265A and S265D mutants could still be visualized by immunoblotting and was not further modulated by RT. We thus validated DNA-PKcs and RAD51 as novel interaction partners of TIPRL1, and concluded that their increased binding upon RT is likely not due to RT-induced TIPRL1 S265 phosphorylation. In contrast to DNA-PKcs, RAD51 plays a major role in HR repair of DSBs (Baumann & West, 1998; Lee & Paull, 2021). Along with the notion that DSBs primarily activate ATM (Curtin, 2023; Shibata & Jeggo, 2021), the discovery of these novel interactors would imply TIPRL1 functioning mainly in the repair of DSBs. Several PP2A-like phosphatases play a role in NHEJ as well as in HR repair. Both PP2A and PP6 have been implicated in DNA-PKcs activity regulation. PP2A C binding to DNA-PKcs resulted in kinase activation by dephosphorylation of DNA- PKcs and Ku70/80, and promotion of functional Ku-DNA-PK complex formation (Douglas et al, 2010; Wang et al, 2009). Likewise, DSB-induced association of DNA-PK with PP6 activated the kinase (Mi et al, 2009; Shen et al, 2011), without impacting DNA-PKcs autophosphorylation at Ser2056 (Douglas et al, 2010). Although we did neither find any modulation of DNA-PKcs Ser2056 phosphorylation in TIPRL1-depleted cells regardless of RT, the increased interaction of TIPRL1 with DNA-PKcs might still serve other roles, such as the modulation of dephosphorylation of other DNA-PKcs phosphosites or of other phosphoproteins in the DNA-PK complex (*e.g*. Ku70/80, Artemis) that are postulated PP2A- (like) phosphatase substrates (Chowdhury et al, 2005). RAD51 is also a phosphoprotein and its Chk1- mediated phosphorylation at Thr309 is essential for full RAD51 activation in HR repair (Peng et al, 2021). Again, despite an increased interaction of TIPRL1 with RAD51 in irradiated cells, RT-induced modulation of Thr309 phosphorylation was not affected by TIPRL1 depletion. Catalytic inhibition of PP2A-like phosphatases by LB-100, as well as loss of specific PP2A B-type subunits, Aα subunit or activator PTPA, have all been shown to inhibit HR and sensitize cells for DSBs (Ambjørn et al, 2021; Aouida et al, 2019; Kalev et al, 2012; Wei et al, 2013). As our data also suggested decreased repair and radiosensitization upon TIPRL1 depletion, this does again not seem to align well with a phosphatase inhibitory role of TIPRL1, although a recent publication demonstrated sensitization to PARP inhibitors by PP2A activating compounds (SMAPs) via a decrease in HR that was found dependent on RAD51 dysregulation (Avelar et al, 2023). Finally, the contribution of TIPRL1 to DSB repair may also involve its modulation of PP4 (Feng et al, 2016; Gingras et al, 2005), which has been shown to dephosphorylate a large number of DDR regulators (e.g. 53BP1, RPA2) in mammalian cells to promote DNA repair (Chowdhury et al, 2008; Lee et al, 2014, 2012, 2010; Lee et al, 2015; Zheng et al, 2019).

The most ‘hidden’ of the novel interaction partners of TIPRL1 in our interactome analysis appeared to be the nucleosomal histones (H2A, H2B, H3 and H4): their regulated binding to TIPRL1 became only apparent in irradiated cells upon omission of PPIs, or upon Ser265 substitution for Ala or Asp. As histones undergo a large number of post-translational modifications in response to DNA damage (Kim et al, 2019), we propose a model, consistent with our MS data, in which a RT-induced modification of the histones at a DSB specifically promotes binding to non-phosphorylated TIPRL1, and thereby localizes it in the immediate vicinity of activated ATM (**Fig EV4A**). Once phosphorylated by ATM, interaction with the modified histones is disfavored. When Ser265 is substituted for alanine, TIPRL1 cannot be phosphorylated anymore by ATM, and maintains binding with the modified histones upon irradiation (**Fig EV4B**). While a phosphomimic S265D mutant may already bind non-modified histones, its interaction will be released by RT-induced histone modification(s) (**Fig EV4C**). If and how the PP2A-like phosphatases are also involved in these responses and ‘accompany’ TIPRL1 to these sites of DNA-damage remains unclear, but is possible, since their interaction with TIPRL1 is not modulated by TIPRL1 phosphorylation. In any case, the opposite binding behavior of the histones to TIPRL1 S265A and S265D mutants clearly correlates well with the opposite phenotypes of irradiated TIPRL1 S265A (radiosensitization) and TIPRL1 S265D (radioresistance) expressing cells, inferring an important role of this Ser265 phosphorylation-dependent, regulated TIPRL1-histone binding in mediating TIPRL1’s function(s) in the DDR. As a final, additional conclusion from our AP/MS experiments, we can state that TIPRL1 Ser265 phosphorylation is also at least in part responsible for the significantly decreased number of TIPRL1 interactors found at 4h post-RT in general, as the opposite (i.e. a higher number of interactors) was seen in the absence of PPIs (i.e. under conditions in which RT-induced Ser265 phosphorylation was not preserved).

The mechanisms underlying the role of TIPRL1 in the DDR discovered here may further guide the rational development of radiosensitizing approaches in TIPRL1-high expressing HNSCC and result in the eventual implementation of TIPRL1 as a biomarker in the clinic. However, more work is required to provide additional mechanistic insights into the precise role of TIPRL1 in the DDR, particularly in the different DSB DNA repair pathways (NHEJ and HR). Based on our current findings, ATM inhibitors, which would inhibit TIPRL1 Ser265 phosphorylation in irradiated cells, might have the highest theoretical radiosensitizing potential, and several clinical trials with these inhibitors in solid cancers are currently underway (Curtin, 2023). Combination therapy with ATM inhibition and RT has already been achieved in solid tumors (Waqar et al, 2022), although potential side-effects are a concern (Weber & Ryan, 2015). Alternatively, PP2A-targeted compounds that might more directly affect the PP2A-TIPRL1 interaction (Arribas et al, 2020, 2023), or new compounds interfering with TIPRL1- RAD51 or TIPRL1-DNA-PKcs binding could become valuable alternatives for testing. Along a different line of thinking, TIPRL1 inhibition in combination with Chk1 inhibitors may also have a therapeutic potential, as inhibition of Chk1-dependent checkpoint maintenance may prove synthetically lethal in TIPRL1-inhibited cells. Although these therapeutic options are still speculative, our study has at least set a basis to consider TIPRL1 targeting as a radiosensitizing approach in HNSCC.

## Materials and Methods

### Cloning, site directed mutagenesis and recombinant protein production

Guide RNAs (gRNA) targeting TIPRL1 were designed using the Custom Alt-R® CRISPR-Cas9 guide RNA (IDT). Single guide RNAs (sgRNA) were cloned into pSpCas9(BB)-2A-Puro (PX459) V2.0, a gift from Feng Zhang (Addgene plasmid #62988; http://n2t.net/addgene:62988) (Ran et al, 2013). Wild-type (WT) TIPRL1 cDNA was cloned into 3xFLAG pLenti6.4 using In-Fusion® HD Cloning Kit (Takara, #639650). Phosphomimic and non-phospho mutants (S265D and S265A) were generated by site-directed mutagenesis using PWO DNA Polymerase (Roche, #11644955001). Primers containing the appropriate mutations were designed with QuickChange® Primer Design Program (Agilent). All primers (**Appendix Table S1**) were ordered at IDT. WT and S265D mutant TIPRL1 were subcloned into pET15b (Sigma, #69661) for recombinant protein production and metal affinity purification as described (Sents et al, 2017).

### Cell culture and lentiviral transduction

HEK293T, Cal27, SQD9, SC263, SCC61 (all from ATCC), and HTEpiC (Sciencell, #2560) cell lines were kept in culture at 37°C and 5% CO_2_. All cells, except HTEpiCs, were cultured in DMEM (Sigma, #D6546) supplemented with 10% fetal bovine serum (FBS; Sigma, #F7524 & Biowest, #S1600), 2mM L-glutamine (Sigma, #G7513) and penicillin/streptomycin (100 units/mL and 10 mg/mL, respectively) (Sigma, #P0781). HTEpiCs were kept in Poly-L-Lysine (2 µg/cm²; Sigma, #P4707) coated culture flasks in TEpiCM-b medium (#2561-b) supplemented with TEpiCGS (#2572) and 5 mL penicillin/streptomycin (#0503) (all from Sciencell). All experiments were performed with *Mycoplasma*-free cells (regularly tested with Venor™ GeM, Minerva Biolabs, #11-1050). HEK293T cells were transfected with TurboFect™ (Thermo Fisher, #R0531) for lentivirus production, followed by transduction of SQD9 cells as described (Meeusen et al, 2021).

### Radiotherapy and cell treatments

SQD9 cells were exposed to 6 Gray (Gy), unless specified otherwise, using a SARRP device (Xstrahl Medical & Life Sciences). RT was delivered using 220 kV photons, 13 mA, a dose rate of 4 Gy/min with a source-to-skin distance of 35 cm and a broad focus. Kinase inhibitors and other DNA damagers were all from Selleck Chem. Stock solutions of the kinase inhibitors (AZD6738 (#S7693), AZD1390 (#S8680), AZD7762 (#S1532) and AZD7648 (#S8843)) of 10 mM were prepared in dimethylsulfoxide (DMSO) (Sigma, #472301), as was etoposide (#1225) and veliparib (ABT888, #S1004, 50 mM). Stock solutions of 1.67 mM cisplatin (#S1166) and 10 mM gemcitabine (#S1149) were prepared in 0.9% NaCl.

### CRISPR/Cas9 gene editing

SQD9 cells were transfected with PX459 using Nucleofector™ 2b Device and Cell Line Nucleofector™ Kit L (Lonza, #AAB-1001, #VACA-1005). Cas9-expressing cells were selected with 0.625 µg/mL puromycin (Thermo Fisher, #A113803) and single cells were allowed to grow into colonies for 2-3 weeks. Colonies were transferred to 96-well plates and grown until a sufficient number of cells was reached for genomic DNA (DNeasy Blood & Tissue Kit; Qiagen, #69504) and protein extraction. Cas-OFFinder (CRISPR RGEN Tools) was used to screen the human genome for potential sgRNA off-targets. Screening of positive clones and off-target analysis on DNA level were based on PCR amplification of the region of interest, followed by Sanger sequencing (LGC genomics) (**Appendix Table S1**).

### 2D colony growth assay

SQD9 cells were seeded in 6-well plates to adhere overnight. The next day, they were irradiated with 0, 2, 4 or 6 Gy, counted and reseeded in 10 cm plates. After 2- to 3-week-incubation, cells were fixed and stained with 0.1% crystal violet in 20% methanol. Colonies containing at least 50 cells were selected for further analysis with Gelcount (Oxford Optronix). Plating efficiencies (PEs) were calculated by dividing the number of colonies by the number of cells seeded. Survival fractions (SFs) were calculated by dividing the PEs of the corresponding radiation dose by the PE for 0 Gy. To obtain dose enhancement factors (DEFs), ratios between the SFs of TIPRL1 WT or S265D expressing cells and those of *TIPRL* indel cells or TIPRL1 S265A expressing cells were calculated for each irradiation dose.

### Protein extraction and western blotting

Cells were collected using cell scrapers, washed twice with ice-cold phosphate-buffered saline (PBS) and lysed in RIPA (50 mM Tris-HCl [pH 7.4], 1% NP-40, 150 mM NaCl, 0.1% sodium deoxycholate, 0.1% SDS, 1 mM EGTA and 1 mM EDTA) supplemented with cOmplete™ protease inhibitors (Roche, #4693116001) and PhosSTOP™ phosphatase inhibitors (Roche, #4906845001), unless otherwise specified. Lysates were incubated on ice for 10 min, vortexed three times and centrifuged (13000 g, 4°C, 15 min). To ensure equal loading, Pierce™ BCA (Invitrogen, #23225) was performed to determine protein concentration before diluting the lysates in sample buffer (4X NuPAGE™ LBS) (Invitrogen, #NP007) and boiling (10 min, 96°C). Total lysates (10-20 µg) were separated by SDS-PAGE on NuPAGE™ 4-12% Bis-Tris Midi Protein Gels (Invitrogen) or 3-8% Tris-Acetate Criterion^TM^ XT Precast Gels (Bio-Rad Laboratories, Invitrogen) at 150 V for 90 min. Proteins were transferred to nitrocellulose (Cytiva, #10600002) or PVDF membrane (Cytiva, #10600023) by wet blotting (40-60 min, 100 V). Membranes were stained 10 min in Ponceau (Thermo Fisher, #BP103), and blocked for 1h at room temperature in 5% non-fat dry milk in Tris-buffered saline (TBS)-Tween®20 (0.1%) (TBS-T). Membranes were incubated overnight at 4°C with primary antibodies (**Appendix Table S5**), washed four times for 15 min with TBS-T, incubated with horseradish peroxidase (HRP)-conjugated secondary antibodies (**Appendix Table S5**) for 1h at room temperature, and washed four times with TBS-T for 10 min. Protein bands were visualized with WesternBright® HRP ECL substrate (Advansta, #K-12045- D50) using ImageQuant LAS 4000 imager (GE Healthcare). Densitometric analysis was executed with Image Studio™ Lite (LiCOR).

### MS-based (phospho) interactomics and targeted phospho analysis of affinity purified (AP) TIPRL1

For AP-MS, protein lysates of TIPRL1_WT cells (no FLAG-tagged TIPRL1) or Rescue TIPRL1_WT/S265A/S265D cells (stably expressing a 3xFLAG-tagged TIPRL1 variant) were incubated for 2h at 4°C with 15 µL anti-FLAG conjugated Sepharose beads (ANTI-FLAG® M2 affinity gel, Sigma, A2220) in 500 µL NENT-100 (20 mM Tris HCl [pH 7.4], 1 mM EDTA, 0.1% NP-40, 25% glycerol and 100 mM NaCl) containing 1 mg/mL bovine serum albumin (BSA), supplemented with cOmpleteTM protease inhibitor and PhosSTOP^TM^ phosphatase inhibitor cocktail, unless specified otherwise. Beads were extensively washed: twice with 500 µL NENT-100, and twice with NENT-300 (20 mM Tris HCl [pH7.4], 1 mM EDTA, 0.1% NP-40, 25% glycerol and 300 mM NaCl). For unbiased (phospho) interactomics, beads were additionally washed twice with 50 mM Tris HCl (pH 8.5) containing 50 mM NaCl, twice with Tris HCl (pH 8.5) containing 0.0025% ProteaseMAX (Promega, #V2072), and subjected to overnight on-bead digestion in the presence of 50 mM Tris HCl (pH 8.5) containing 5% CH_3_CN, 0.01% ProteaseMAX and 0.5 µg trypsin (Thermo Fisher, #90058). The resulting peptide mixture was desalted on C18 spin columns (Thermo Fisher, #89870). For unbiased phospho analysis of the immunoprecipitates, peptides were desalted on Macro Spin Tips (Harvard Apparatus) before processing on Phos-TiO_2_ tips (GL Sciences, #GL-5010-21308) essentially according to the manufacturer’s instructions, but in the presence of 2.5% lactic acid. For targeted TIPRL1 phospho analysis, the beads were supplemented with 20 µL sample buffer, boiled for 10 min and resolved on SDS-PAGE. Gels were stained for 3h with Colloidal Coomassie (Invitrogen, #LC6025) and destained overnight in MQ water. The TIPRL1 band was excised and digested with 0.3 µg trypsin in the presence of 200 mM NH_4_HCO_3_ and 0.01% ProteaseMAX. After absorption, 30 µl of the solution was added to the gel pieces and left overnight at 37°C. After elution, peptides were desalted by C18 ZipTips (Millipore, #ZTC18S096).

LC-MS/MS analysis was executed on an Ultimate 3000 UPLC system (Dionex, Thermo Fisher) equipped with an Acclaim PepMap100 pre-column (C18, particle size 3 μm, pore size 100 Å, diameter 75 μm, length 20 mm) and a C18 PepMap analytical column (particle size 2 μm, pore size 100 Å, diameter 50 μm, length 150 mm) using a 40 min linear gradient (300 nL/min) coupled to a Q Exactive Orbitrap mass spectrometer (Thermo Fisher) operated in data-dependent (phospho interactomics) or tSIM-ddMS^2^ (targeted TIPRL1 phospho analysis) acquisition mode. For unbiased (phospho) interactomics of the immunoprecipitates, peptides were identified by Mascot (Matrix Science) through Proteome Discoverer 2.2 (Thermo Fisher) using Uniprot *Homo sapiens* database (204906 entries). Oxidation (M) (and Phospho S/T if applicable) were included as variable modifications. Two (three for phospho analysis) missed cleavages were allowed, peptide tolerance was set at 10 ppm for MS and at 20 mmu for MS/MS. Proteome Discoverer PhosphoRS node was used for localizing the phosphorylation. Peptides were validated using Scaffold software (Protein FDR 1%, Peptide FDR 1%, minimum 1 peptide). Progenesis (Nonlinear Dynamics) was used for relative quantification of peptides. Only exclusive peptides were used for protein quantification.

Proteins were considered as TIPRL1 interactors if they were (1) at least 1.87 (the cut-off ratio was determined based on a frequency distribution of the log of all enrichment factors) times enriched, after normalization to the sum of all feature abundances, not taking into account bait feature abundances, in the considered condition versus the ‘no FLAG control’, and (2) if the spectral counts of the protein in that condition (obtained from Scaffold) were above zero. To gauge differential binding, protein abundances were normalized to the bait abundance in each condition. The cut-off ratio for differential binding was determined based on a frequency distribution of the log of all protein ratios between two conditions. For targeted phospho analysis, data were processed through Skyline software. The mono-isotopic peak area of each phosphorylated TIPRL1 peptide in the different conditions was normalized to the average mono-isotopic peak area of non-phosphorylated TIPRL1 peptides in each condition.

### MS-based proteomics analysis of TIPRL1_WT and TIPRL1 indel_2 cells

Cell pellets were washed in TBS, and lysed in 4% (w/v) sodium deoxycholate (SDC) in 50 mM Tris HCl (pH 8.5) (Humphrey et al, 2018) to obtain a protein concentration of 7-11 mg/ml. Samples were promptly heat-inactivated upon lysis (10 min, 96°C) and subjected to sonication on ice for 3 times 30s cycles (1s on and 1s off). One mg of each sample was subjected to reduction, alkylation and quenching in the presence of 10 mM dithiothreitol (DTT), 25 mM iodoacetamide and 25 mM DTT, respectively. Samples were subsequently diluted five times in Tris HCl (pH 8.5) containing 10 µg trypsin (Thermo Fisher, #90058).

For unbiased proteomics analysis, SDC was removed by adding 0.5% TFA followed by centrifugation (10 min at 14,000 g). Exactly 83% of the supernatant was made up to 100 µl with water, brought to 1% TFA and again centrifuged (10 min at 14,000 g). Eighty µl was loaded on C18 Pierce desalting columns (Thermo Fisher, #89870) and eluted with 60% CH_3_CN, 0.1% TFA. One tenth of the sample (2 µg) was analyzed by LC-MS/MS on a Q Exactive orbitrap (see above) using an LC column of 500 mm length. Protein lists were obtained using Progenesis (Nonlinear Dynamics) label-free quantification software, with quantifications only considering peptides with a Mascot score higher than 29 (95% probability threshold) and including all peptides per protein. Normalized protein abundances were subjected to rank regression (p<0.05) as a function of time after 4h of radiotherapy (Qlucore software). The resulting output list was loaded into David (Sherman et al, 2022) using the total unbiased protein list (3793 entries) as a background. Enriched clusters were retrieved using the ‘Functional Annotation clustering’ tool (high stringency and EASE score <0.05). Heat maps were drawn using Qlucore software.

### Immunofluorescence

Cells were seeded in 96-well Black/Clear bottom plates. The next day, they were irradiated (2 or 6 Gy), and fixed with 4% paraformaldehyde at different time points after irradiation. Using Triton X-100 (Sigma, #T8787) cells were made permeable for P-Histone H2A.X (Ser139) (Millipore, cloneJBW301) or DYKDDDDK (FLAG) Tag Monoclonal Antibody (Sigma, #F1804). Nuclei were counterstained with DAPI (Sigma, #D9542) and imaged using Operetta CLS (Perkin Elmer). Analysis was performed with ImageJ. For immunofluorescence analysis with the FLAG antibody (n=1), six technical replicates were counted.

### Flow cytometry

Flow cytometry analyses were executed on a FACSverse device (BD Biosciences). For cell cycle analysis, cells were irradiated with 6 Gy and DNA was labeled with 75 µM Bromodeoxyuridine (BrdU) (Sigma, #B5002) 1h before the indicated time points. Another hour after the indicated times, cells were fixed with 70% ethanol (Thermo Fisher, #397690010). DNA of the fixed cells was denatured in 2N HCl (Chem-Lab, #CL00.0310)/Triton X-100, and the solution neutralized with 0.1M Na_2_B_4_O_7_ (Sigma, #221732), pH8.5. Next, 20 µL Anti-BrdU-FITC (Becton Dickinson, #347583) per 10^6^ cells was added and cells were stained with 10 µg/mL propidium iodide (PI) (Sigma, #P4170) supplemented with 100 µg/mL RNase-A (Invitrogen, #12091021). For cell death analysis, cells were collected at indicated time points, and stained with PI supplemented with Annexin V from the Annexin-V-FLUOS Staining kit (Roche, #11858777001).

### In vitro PP2A activity assay

PP2A dimer was purchased from SignalChem (PP2Aα/PPP2R1A Active Complex, referred to as PP2AD). *In vitro* phosphatase activity was measured on K-R-pT-I-R-R phosphopeptide (SynPeptide). PP2AD was preincubated for 15 min at 30°C in 20 mM Tris/HCl (pH7.4) with recombinant TIPRL1 WT or S265D mutant, purified from bacteria as described (Haesen et al, 2016). The phosphatase reaction (20 µl) was initiated by adding 9 µl substrate (2 mM). Reactions were stopped after 7.5, 15 and 30 min by adding BIOMOL green (Enzo Life Sciences, #BML-AK111). Absorbance at 620 nm was measured in a multi-channel spectrophotometer another 30 min later.

### Immunohistochemistry

Paraffin-embedded HNSCC tissue sections were obtained from a patient cohort available in UZ Leuven, following initial surgery (study approval: S54731 (Deschuymer et al, 2020)). Sections were dried, and hematoxylin/eosin staining was performed using an automated procedure (Leica Microsystems). Immunohistochemical (IHC) staining with TIPRL1 antibody (1:200; Abcam, #ab70795) was conducted on an automated IHC system (Bond™ Max, Leica Biosystems). A 3,3′- diaminobenzidine solution (Liquid DAB^+^ Substrate Chromogen System, Dako) was used as chromogen. Slides were counterstained with hematoxylin and mounted with a CV5030 Glass Coverslipper (Leica Biosystems). Pictures were acquired on a DM2000 Histology Microscope (Leica Microsystems). Tumors were classified into two groups according to low and high TIPRL1 expression, with high expression defined as >25% of tumor cells showing positive staining.

### cBioportal analysis

Using the HNSCC TCGA (Firehose Legacy) study in the cBioportal database (Cerami et al, 2012; Gao et al, 2013; The Cancer Genome Atlas Network, 2015), patients were stratified based on their HPV status, treatment (RT or no RT) and TIPRL1 status. For the latter, mean tumoral TIPRL1 expression was calculated, and two groups were formed: one with low and one with high expression (relative to the mean); TIPRL1-high and TIPRL1-low tumors were subsequently correlated with patient survival data using RStudio.

### Statistics and data availability

Statistical analysis was executed in Graphpad or RStudio using the appropriate statistical tests (detailed in figure legends). Statistical differences are indicated with *: 0.01< p <0.05; **: 0.001< p <0.01; and ***: p <0.001.

All mass spectrometry interactomics and proteomics data have been deposited to PRIDE. Source data (uncropped blots, raw data values of graphs with multiple replicates, …) can be made available.

## Acknowledgments and funding

We would like to thank H. Geeraerts and L. Lenaerts for technical assistance in the irradiation experiments and MS sample preparations. Prof. P. Agostinis (CDRT Laboratory, KU Leuven) is acknowledged for use of the Amaxa Nucleofector device. Several experiments were performed at KU Leuven core facilities (SyBioMA, MOSAIC and Bioimaging Core). A. Sablina, S. Nuyts, R. Derua and V. Janssens were jointly funded by a KU Leuven C2-project (C24/17/073).

## Author contributions and conflicts of interest

Conceptualization: CC, RüD, AS, SN, RD, VJ; Data acquisition and analysis: CC, RüD, EEC, PZ, RD; Study supervision: RD, VJ; Funding acquisition: AS, SN, RD, VJ; Writing – original draft: CC, RD, VJ; Writing – review & editing: all authors.

None of the authors have any competing interests to declare.

## Expanded View Figure Legends

**Figure EV1.**
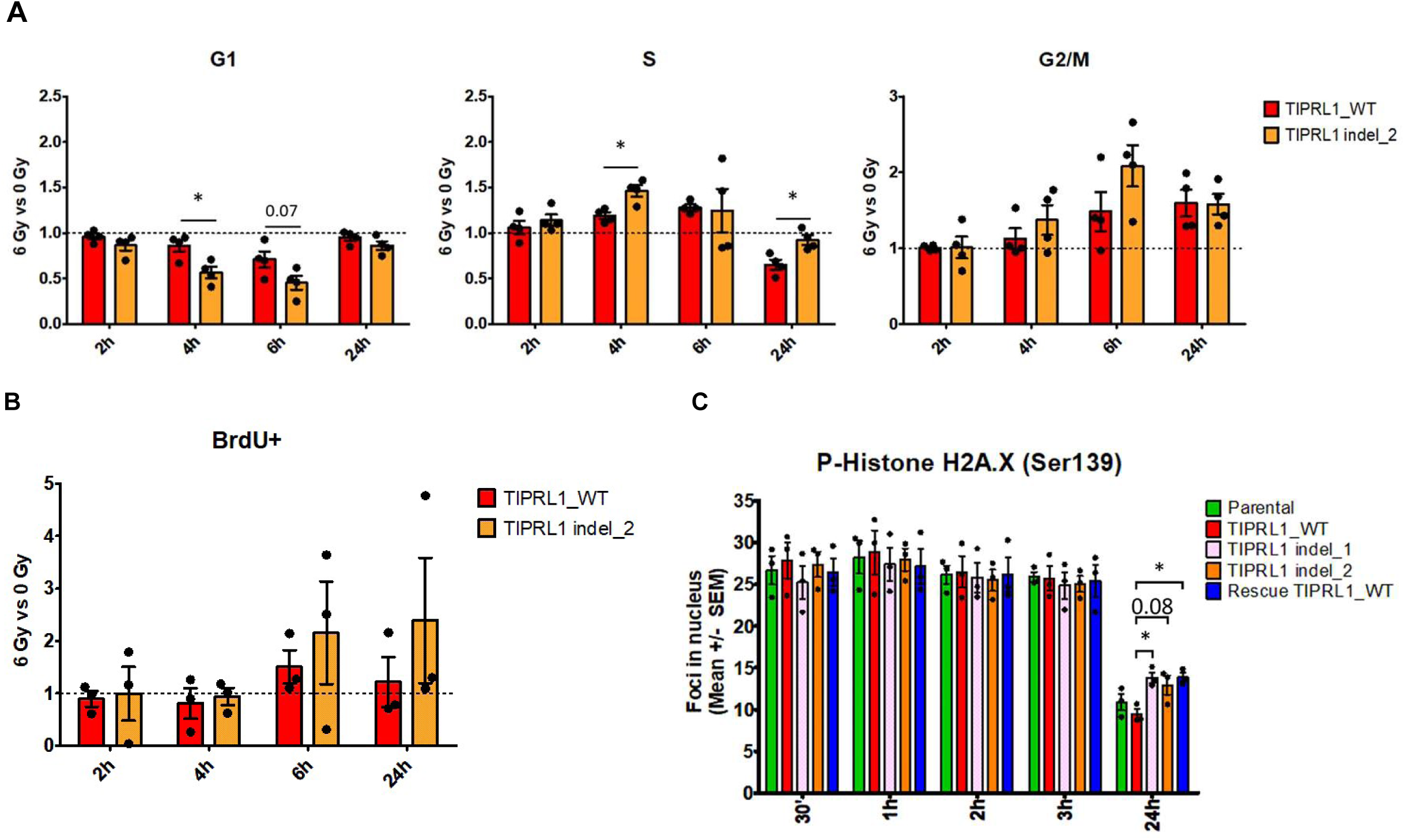
TIPRL1 indel cells show faster and prolonged S phase checkpoint arrest after DNA damage. **A** Cell cycle distribution as measured by PI staining and flow cytometry. For each cell cycle phase the ratio of cells in this phase with irradiation (6 Gy) was calculated versus without irradiation. T-test was used for analysis between TIPRL1 indel_2 and TIPRL1_WT SQD9 cell lines per time point after irradiation (n=4, mean ± SEM, *: p<0.05). **B** Fraction of cells undergone S-phase as measured by BrdU staining and flow cytometry. Ratios were calculated of BrdU-positive cells with 6 Gy vs 0 Gy for each time point and both cell lines (n=3, mean ± SEM, t-test). **C** SQD9 cells were exposed to 2 Gy irradiation and fixed for immunofluorescence staining at different time points after treatment to determine Phospho-Histone H2A.X (Ser139) foci formation (n=3, mean ± SEM, one-way ANOVA followed by Tukey’s multiple comparison test, *: p<0.05).

**Figure EV2.**
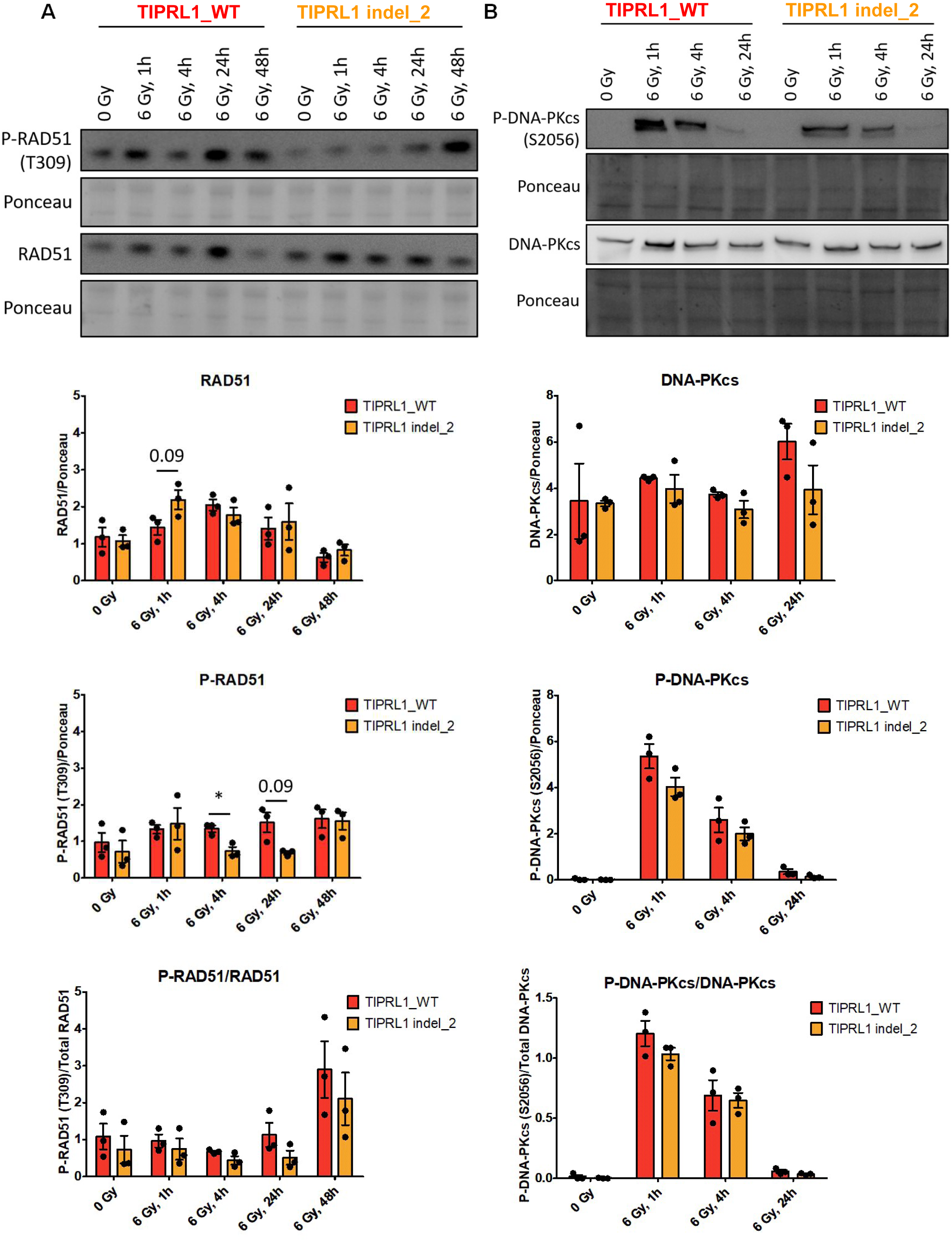
Loss of TIPRL1 expression does not affect RT-induced RAD51 T309 and DNA-PKcs S2056 phosphorylation. **A – B** TIPRL1_WT and TIPRL1 indel_2 cells were irradiated (6 Gy), and protein lysates were made at several times post-RT (0 Gy, 1h, 4h, 24h and 48h) for western blot analysis. Membranes were developed with primary RAD51 and P-RAD51 (Thr309) antibodies (**A**), and primary DNA-PKcs and P- DNA-PKcs (S2056) antibodies (**B**). Ponceau was used as loading control. Quantifications were performed with Image Studio™ Lite software and present mean values ± SEM (n=3, t-test, *: p<0.05).

**Figure EV3.**
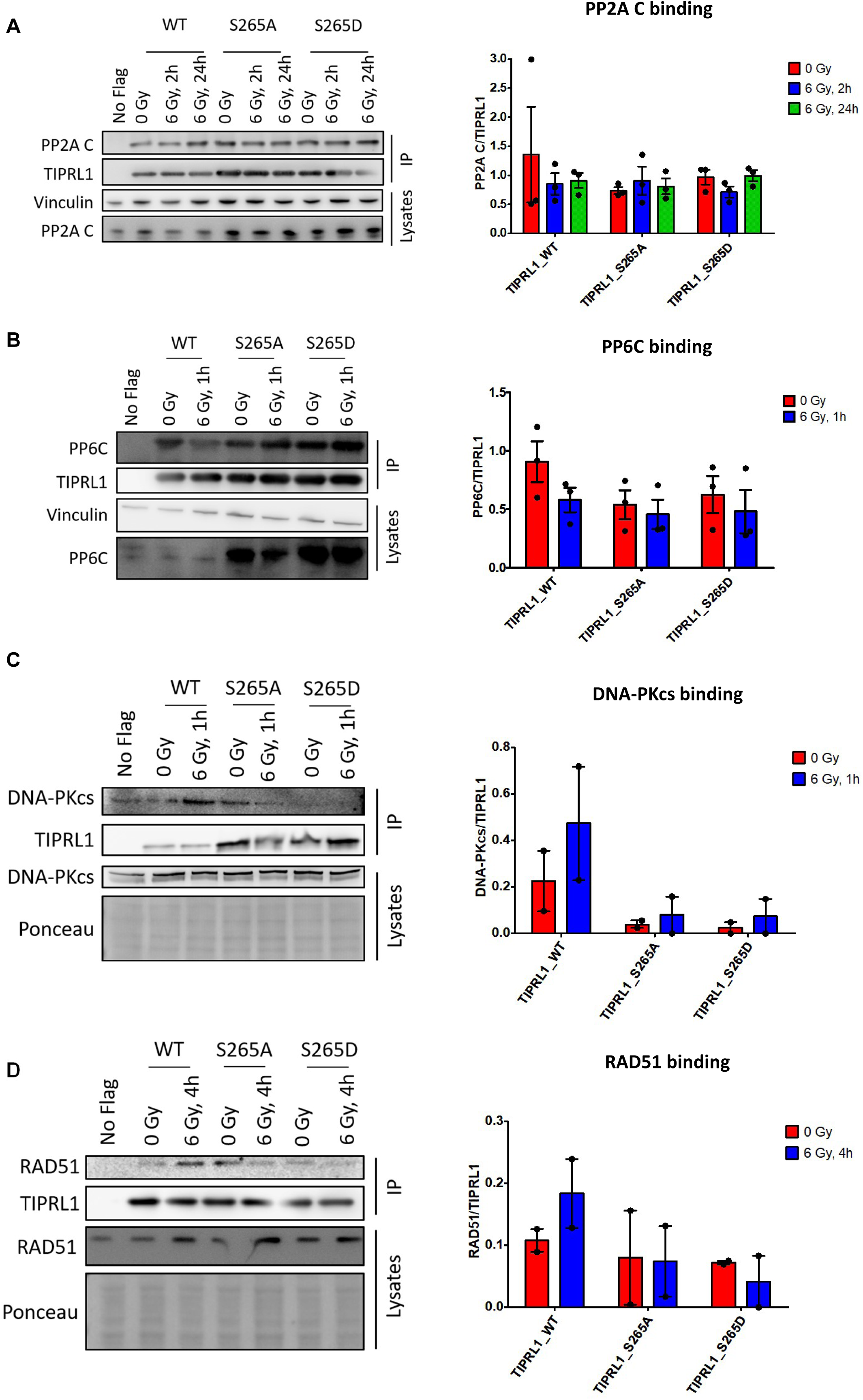
Effects of irradiation-induced TIPRL1 phosphorylation on PP2A-like phosphatase, DNA-PKcs and RAD51 binding. **A – D** Protein lysates of non-irradiated (0 Gy) or irradiated (6 Gy) TIPRL1_WT (No Flag), Rescue TIPRL1_WT, Rescue TIPRL1_S265A and Rescue TIPRL1_S265D cells were prepared at indicated times post-RT, and subjected to anti-FLAG immunoprecipitation (IP). Representative blots of PP2A C (**A**), PP6C (**B**), DNA-PKcs (**C**) and RAD51 (**D**) expression and retrieval in the IPs in function of irradiation are shown (left panels). Lysate preparation and IPs were performed in the presence of PPIs (to preserve TIPRL1 S265 phosphorylation), except for (**A**) (to allow detection of the PP2A-TIPRL1 interaction). Corresponding quantifications (right panels) of TIPRL1–PP2A C binding (n=3, mean ± SEM, one-way ANOVA followed by Tukey’s multiple comparison) (**A**), TIPRL1–PP6C binding (n=3, mean ± SEM, one-way ANOVA followed by Tukey’s multiple comparison) (**B**), TIPRL1–DNA-PKcs binding (n=2, mean ± SEM) (**C**) and TIPRL1–RAD51 binding (n=2, mean ± SEM) (**D**) are shown.

**Figure EV4.**
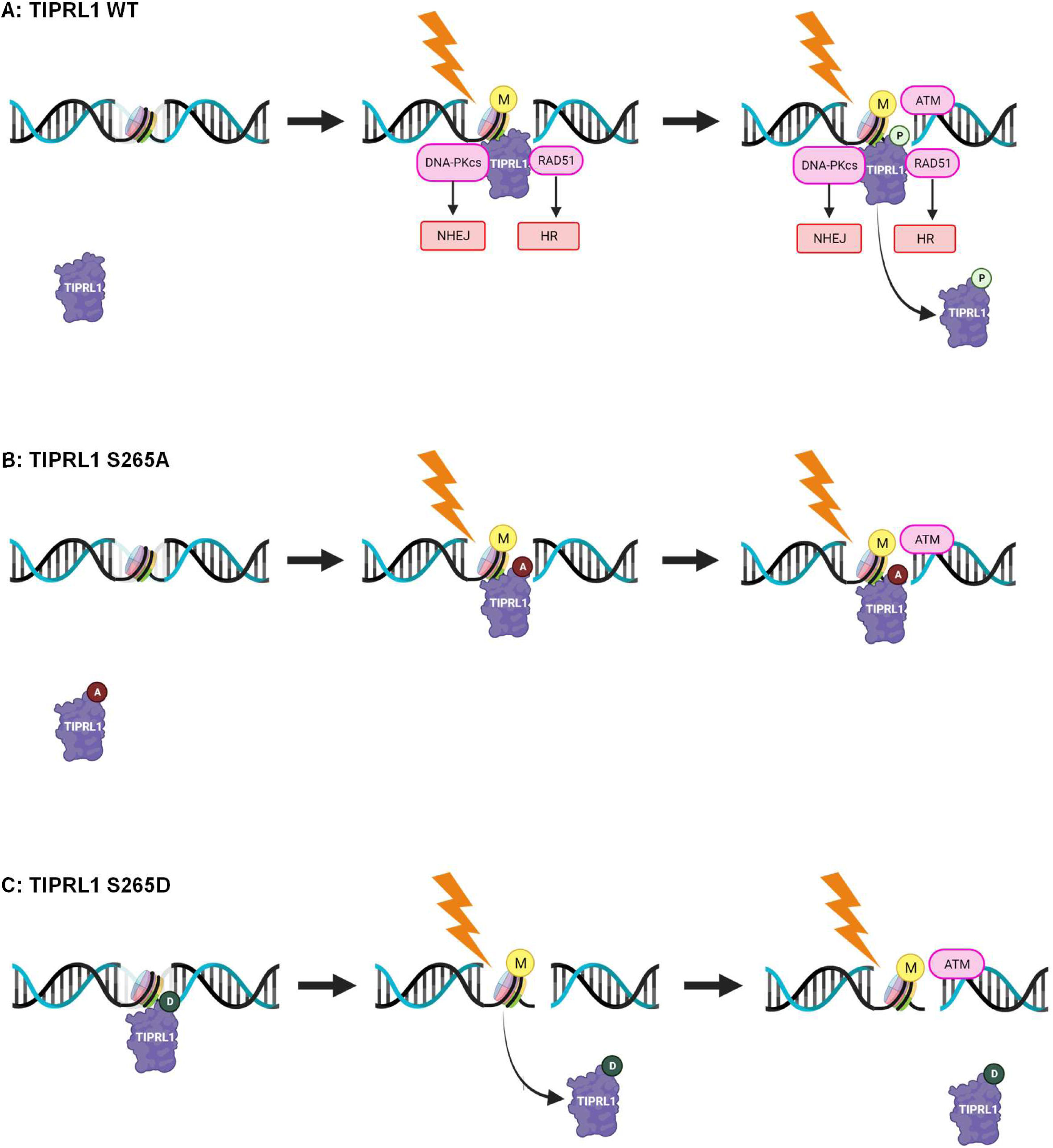
Proposed model on the role of ATM-dependent TIPRL1 phosphorylation upon DNA damage. Hypothesis shown for TIPRL1 WT (**A**), TIPRL1 S265A (**B**) and TIPRL1 S265D (**C**). The model implies two types of modifications that are both induced by RT (or, likely more general, by DSBs): a (not further investigated/specified) histone modification (M) and the newly identified ATM-dependent TIPRL1 phosphorylation at S265 (P). In the absence of both modifications (no RT), TIPRL1 does not bind any histones. In the presence of one of either modification (or D phosphomimic), binding is promoted. In the presence of both modifications, binding is inhibited. The model provides a rationale for TIPRL1 recruitment through (RT-induced, modified) histones in the vicinity of activated ATM, and for subsequent release of phosphorylated TIPRL1 from these sites. TIPRL1 interaction with DSB repair factors, DNA-PKcs (involved in Non-Homologous End Joining, NHEJ) and RAD51 (involved in Homologous Recombination repair, HR) appears not strictly affected by TIPRL1 S265 phosphorylation, but is increased by RT. If and how TIPRL1 might regulate the functions of the latter to promote DNA repair, and whether that involves inhibition of PP2A-like phosphatases PP2A, PP4 and/or PP6, remains an open question.

